# Spatially Tuned Localization of Interleukins and OX40 Agonists Enhances Synergistic Anti-Tumor Immunity

**DOI:** 10.1101/2025.05.15.654266

**Authors:** John H. Klich, Anahita Nejatfard, Emily L. Meany, Ben S. Ou, Julie Baillet, Eric A. Appel

## Abstract

Advances in immunotherapy have revolutionized the current standard of care for cancer patients, but unfortunately, most approaches still fail to mount a robust anti-cancer effect. This is in part due to a highly immunosuppressive tumor microenvironment which has developed bio-orthogonal mechanisms of immune escape. To address this challenge, the field has turned to combination immunotherapies, but systemic administration of potent combination therapies has resulted in severe immune related adverse effects and toxicity. Enabling potent combination immunotherapies requires administering these immune agonists in a way that more closely resembles the endogenous cancer immunity cycle, a tightly regulated sequence of cues in both space and time. Here, we explore the ability of an injectable hydrogel depot to enable the rational localization of potent immunotherapeutic cytokines (IL-12, IL-2) and antibodies (OX40a). We hypothesized that selectively altering the biodistribution of these cargo would enable tolerable and synergic anti-cancer combinations, so we leveraged a previously characterized injectable polymer-nanoparticle (PNP) hydrogel system to deliver these agonists either intratumorally (IT) or peritumorally (PT). Using *in vivo* imaging, we demonstrated that site of administration is critical to redistributing cargo to either the tumor or tumor draining lymph node (tdLN). Further, we demonstrated that the targeted localization of cytokine and antibody therapies synergistically improved treatment efficacy in the B16F10 and MC38 tumor models and altered cellular phenotypes in these microenvironments. This approach thus represents a crucial new strategy for basic cancer immunology and materials-based immuno-engineering research while improving therapeutic efficacy.

## 1. Introduction

Despite the incredible progress made in developing novel cancer immunotherapies, cancer remains the second leading cause of death in the United States in 2024, with projections of over two million new diagnoses and over 600,000 cancer fatalities (*1*). Even more, the incidence of six of the top ten cancers is increasing. And these trends disproportionately affect countries with lower development indices, indicating the need for broadly translatable next generation immunotherapies (*1, 2*).

Immune checkpoint blockade therapy has been adopted as a frontline cancer immunotherapy for improving progression-free survival in patients, (*3–6*) but unfortunately, clinical trials report that up to 75% of patients still fail to respond to treatment (*7, 8*). These results are likely due to a highly immunosuppressive tumor microenvironment (TME) that has evolved multiple, bio-orthogonal mechanisms of immunosuppression, (*9*) indicating the need for combination immunotherapies that target both innate and adaptive axes of anti-cancer immunity.

For the former, immunostimulatory cytokines like interleukin-12 (IL-12) and interleukin-2 (IL-2) have emerged as promising agonists for their complementary activities inducing interferon-γ (IFNγ) mediated T-cell cytotoxicity and stimulating natural killer (NK) cells, repolarizing the tumor microenvironment from a “cold” immune excluded state to a “hot” anti-cancer state (*10*). Despite their potential, little progress has been made in broadly translating these therapies since the approval of IL-2 for melanoma over 30 years ago. The challenge with many immune agonists is their exceptionally small therapeutic window as many cytokines are pleiotropic, resulting in global immune stimulation when delivered systemically. This on-target (correct receptor) / off-tumor (wrong tissue localization) sink drives the necessity for high doses to achieve a therapeutic effect, inadvertently producing immune related adverse effects and dose-limiting toxicities (*11, 12*).

These limitations are symptomatic of the way in which immune agonists are delivered, typically via systemic intravenous infusion. To address these localization related challenges, the field has turned to intratumoral (IT) administration of potent immune agonists, leveraging advancements in radiology, laparoscopy and endoscopy that render many solid tumors accessible for minimally invasive injection (*13*). IT administration alone, however, is not without its limitations. In one clinical trial of IT administered IL-12, peak plasma levels of IL-12 were observed only 5-7 hours post injection, (*14*) suggesting that IT administration needs to be coupled with an effective retention strategy, such as anchoring to the tumor via conjugation of the cargo to tumor-binding peptides or sustained delivery from a biomaterial scaffold. These strategies have been shown to capitalize on the reduced toxicity and improved therapeutic efficacy possible with IT localization.

Several distinct biomaterials approaches conferring sustained, localized IT delivery of immune agonists have been explored. For instance, intratumorally injected alum-tethered cytokines were shown to fundamentally alter their pharmacokinetics, resulting in decreased systemic toxicity while conferring a substantial benefit in treatment efficacy (*15, 16*). In other applications, tumor features like the high deposition of collagen in the tumor extracellular matrix have been used to target and tether cytokines or immune agonists, such as the Toll-like receptor 9 agonist CpG, resulting in significant benefits in safety profiles and treatment potencies (*17–19*). Yet, many of these approaches require modifying the biologic of interest, limiting the scope of cargo that could be administered, and repeat administrations, a process that reduces translatability. To overcome these limitations, hydrogels (water swollen polymer networks with favorable biocompatibility, degradability, and mechanical properties) have shown great promise in improving the safety and efficacy of the physico-chemically distinct biologics often necessary for efficacious combination immunotherapy (*20, 21*).

Further, many strategies have focused on stimulating cytotoxicity but have neglected long-term immunological memory, which is necessary to treat abscopal tumors, metastasis, or recurrences. Next generation immunotherapies should therefore leverage the progress made in IT retention strategies while exploring synergistic combinations of immune agonists that drive both immediate cytotoxicity and immunological memory, the combination of which has been shown to be essential to eradicating large, established immunosuppressive tumors (*22*).

One strategy for mounting a robust memory response is to target immunological processes that occur in the lymph nodes, such as the differentiation and clonal expansion of T lymphocytes upon antigen stimulation and interaction with antigen presenting cells (APCs). Among the T-cell co-stimulatory receptors belonging to the tumor necrosis factor receptor superfamily, OX40 is implicated in critical immunoregulation, including formation of memory T-cell subsets. OX40 agonism has shown promising therapeutic effects both preclinically and in clinical trials as both a monotherapy and a combination therapy (*23–26*).

Here, we investigated how the selective locoregional delivery of distinct immune agonists, enabled by an injectable hydrogel depot technology, could impart unique immunological changes to the tumor microenvironment or the tumor draining lymph node (tdLN), facilitating both cytotoxicity and immunological memory. Based on prior results demonstrating improved safety, tolerability, and potency with the sustained delivery of antibodies and immunostimulatory cytokines, (*21, 27*) we hypothesized that injectable polymer-nanoparticle (PNP) hydrogels could provide sustained site-specific control over the localization of cytokines (IL-12 and IL-2) and agonistic anti-OX40 antibodies to elucidate crucial immunological processes and form the basis for synergistic combination immunotherapy.

We evaluated these immune agonists as monotherapies and combination therapies, delivered either intratumorally or peritumorally, in the poorly immunogenic B16F10 melanoma model, and we validated the lead combination in the MC38 adenocarcinoma model. We examined how site of administration of these therapies skewed biodistribution with *in vivo* imaging of excised organs and explored how this altered biodistribution could be leveraged to enable synergistic combination immunotherapies that drive tumor repolarization and effective immunological memory. We characterized these responses with flow cytometry, demonstrating that site of administration is crucial to mounting distinct immunophenotypes in the tumors and lymph nodes in treated mice. We demonstrated that maximizing the therapeutic efficacy required pro-inflammatory cytokines IL-2 and IL-12 to be delivered intratumorally in combination with peritumorally administered OX40a, where efficacy was substantially diminished if the cargo were co-localized. Notably, this platform approach was demonstrated to be both cargo and tumor agnostic, improving efficacy in multiple tumor models without requiring modification of the cargo, suggesting a promising path for future exploration of novel combination immunotherapies.

## 2. Results

### 2.1. PNP hydrogel depots modulate cargo exposure depending on site of administration

We hypothesized that the sustained, local delivery of immune agonist cargo such as OX40a from injectable hydrogels would increase drug exposure as a function of site of administration. We have previously reported positron emission tomography data demonstrating that hydrogels substantially altered the pharmacokinetics of CD40a to favor drug exposure in target tissues, most notably resulting in retention at the injection site and accumulation in the tdLNs (*27*). We suspected that this feature of sustained locoregional delivery from a PNP hydrogel depot could be exploited to selectively retain cargo in the tumor via IT administration or in the tdLNs via peritumoral (PT) administration (**Fig. 1A**). To this end, we labeled anti-OX40 agonist antibodies (OX40a) with AlexaFluor 647 NHS-ester, according to the manufacturer’s recommendations, and prepared OX40a loaded PNP hydrogels by physically mixing aqueous solutions of dodecyl-modified hydroxypropyl methylcellulose (HPMC-C_12_), poly(ethylene glycol)-poly(lactic acid) (PEG-PLA) nanoparticles, and labeled OX40a. These hydrogels form rapidly upon mixing through dynamic, multivalent, non-covalent interactions between the HPMC-C_12_ polymers and the PEG-b-PLA NPs, enabling facile encapsulation of diverse cargo and easy administration by injection (**Fig. 1A**). PNP hydrogels exhibit robust solid-like properties across a range of physiologically relevant frequencies, shear-thinning behavior necessary for injectability through fine gauge needles used in clinical settings, rapid self-healing that minimizes burst release following injection, and excellent biocompatibility (*28–30*).

**Fig. 1.**
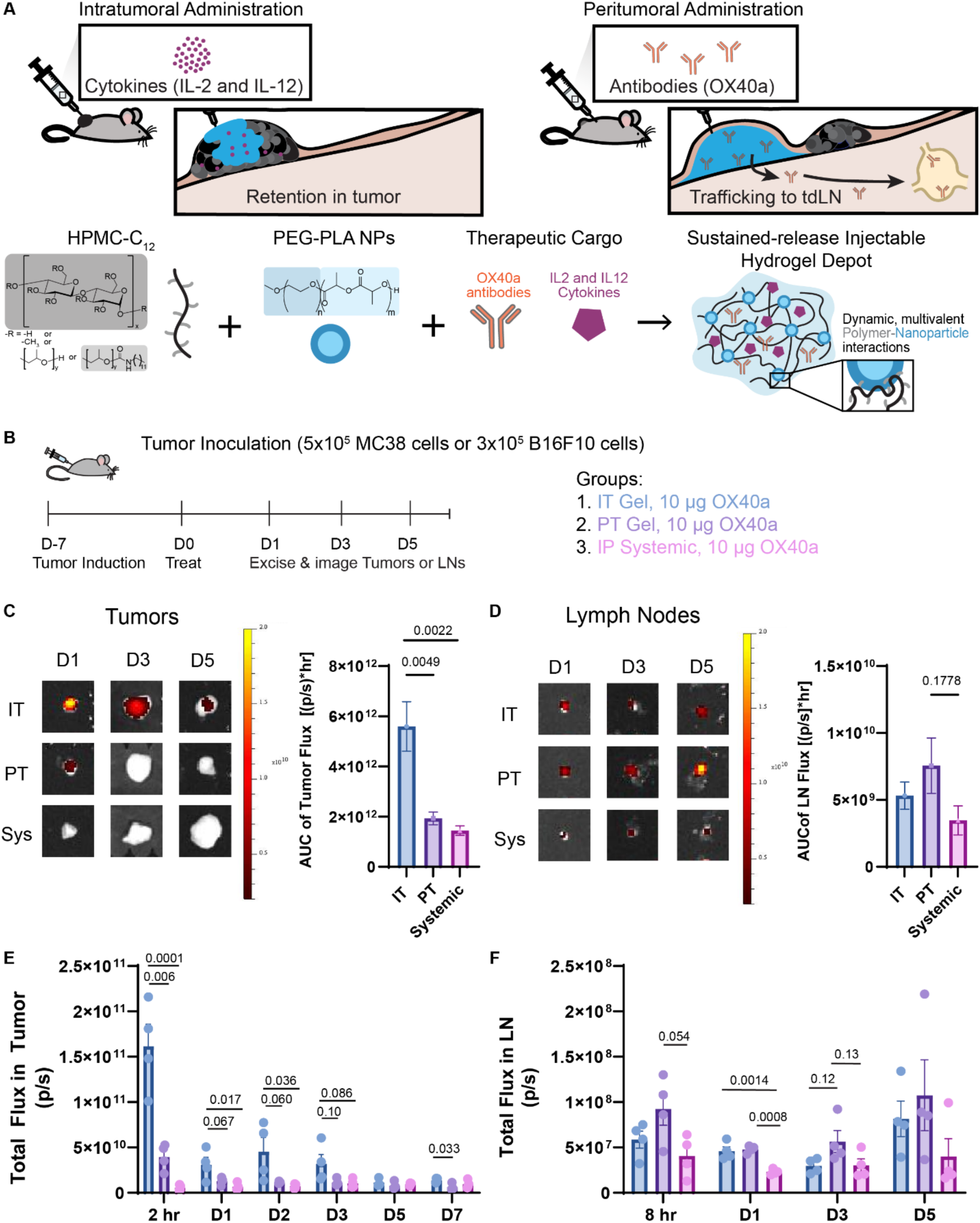
Polymeric nanoparticle hydrogels increase site specific cargo accumulation. (**A**) Schematic of experimental hypothesis indicating that PNP hydrogels can be used to selectively distribute immunotherapies as a function of site of administration, comparing intratumoral and peritumoral injection (top). Schematic of dodecyl-modified hydroxypropyl methylcellulose (HPMC-C_12_) and poly(ethylene glycol)-poly(lactic acid) (PEG-PLA) nanoparticles as functional building blocks of the PNP hydrogel depot technology (bottom). (**B**) Experimental timeline, groups and doses for tumor induction, treatment, and tumor or lymph node excision and imaging by IVIS, with N=4 per group per timepoint. MC38 was used for tumor imaging due to lack of pigmentation while B16F10 was induced in mice from which LNs were excised. (**C**) Representative images of excised tumors on days 1, 3, and 5 from each treatment group (left) and quantification of total exposure via area under the curve of total flux in the tumor over 7 days (right). (**D**) Representative images of excised tumor draining lymph nodes on days 1, 3, and 5 from each treatment group (left) and quantification of total exposure via area under the curve of total flux in the tumor over 5 days (right). (**E**) Total flux in excised tumors for each time point. (**F**) Total flux in excised tumor draining lymph nodes for each time point. Data are reported as mean +/− SEM. Statistics are ordinary one-way ANOVA run in GraphPad Prism with Tukey’s multiple comparisons test.

To evaluate differences in biodistribution and bioaccumulation of cargo from distinct sites of administration, C57BL/6 mice were inoculated with either 5 × 10^5^ MC38 cells or 3×10^5^ B16F10 cells and treated 7 days thereafter either IT or PT with PNP-1-5 hydrogels (*n.b.*, 1-5 refers to the hydrogel formulation comprising 1wt% HPMC-C_12_ and 5wt% PEG-PLA NP, hereafter is referred to simply as PNP hydrogel) loaded with 10 μg of fluorolabeled OX40a. As a clinically relevant systemic control, one group received 10 μg of fluorolabeled OX40a as a soluble intraperitoneal injection (**Fig. 1B**). To assess retention in either the tumor or tdLN, tumors and tdLNs were excised at the timepoints indicated and imaged with an *in vivo* imaging system (IVIS). As an indicator of total bioaccumulation, area under the curve of total flux was quantified over the time series of imaging in both tumors and tdLNs (**Fig. 1C,D**). We found that IT administration resulted in significantly greater retention and accumulation in the tumor compared to either PT or systemic administration of the labeled antibody (**Fig. 1C**). In contrast, PT administration resulted in an increase in antibody exposure to the tdLNs when compared to IT or systemic administration (**Fig. 1D**). The trends in the AUC are consistent with site specific exposure at each time point. Notably, IT administration maintains meaningfully increased tumor localization of cargo through day 7 post treatment while PT administration increases tdLN exposure through day 5 post treatment compared to alternate sites of administration (**Fig. 1E,F**) Collectively, these data demonstrate that the PNP hydrogel depot technology can be leveraged to selectively skew cargo accumulation and total exposure as a function of site of administration. Interestingly, we also noted by inspection meaningful differences in tumor burden depending on site of administration, with those mice receiving PT OX40a having the smallest tumors (**Fig. S1**).

### 2.2 Intratumoral hydrogel administration of cytokines confers survival benefit over peritumoral administration

Having demonstrated unique pharmacokinetics and biodistribution from IT and PT hydrogel depot administration, as well as systemic administration in a standard liquid vehicle, we set out to examine whether these differences resulted in changes in therapeutic efficacy when delivering potent proinflammatory cytokines. First, we evaluated the effect and tolerability of a single administration of low dose IL-12 in C57B/6 mice induced with 3×10^5^ B16F10 cells and treated 4 days thereafter (**Fig. 2A**). Treatment groups included PNP hydrogels loaded with 5 μg of IL-12 administered either IT or PT, empty PNP hydrogel delivered IT as an untreated control, or IT soluble IL-12 as a standard liquid vehicle control.

**Fig. 2.**
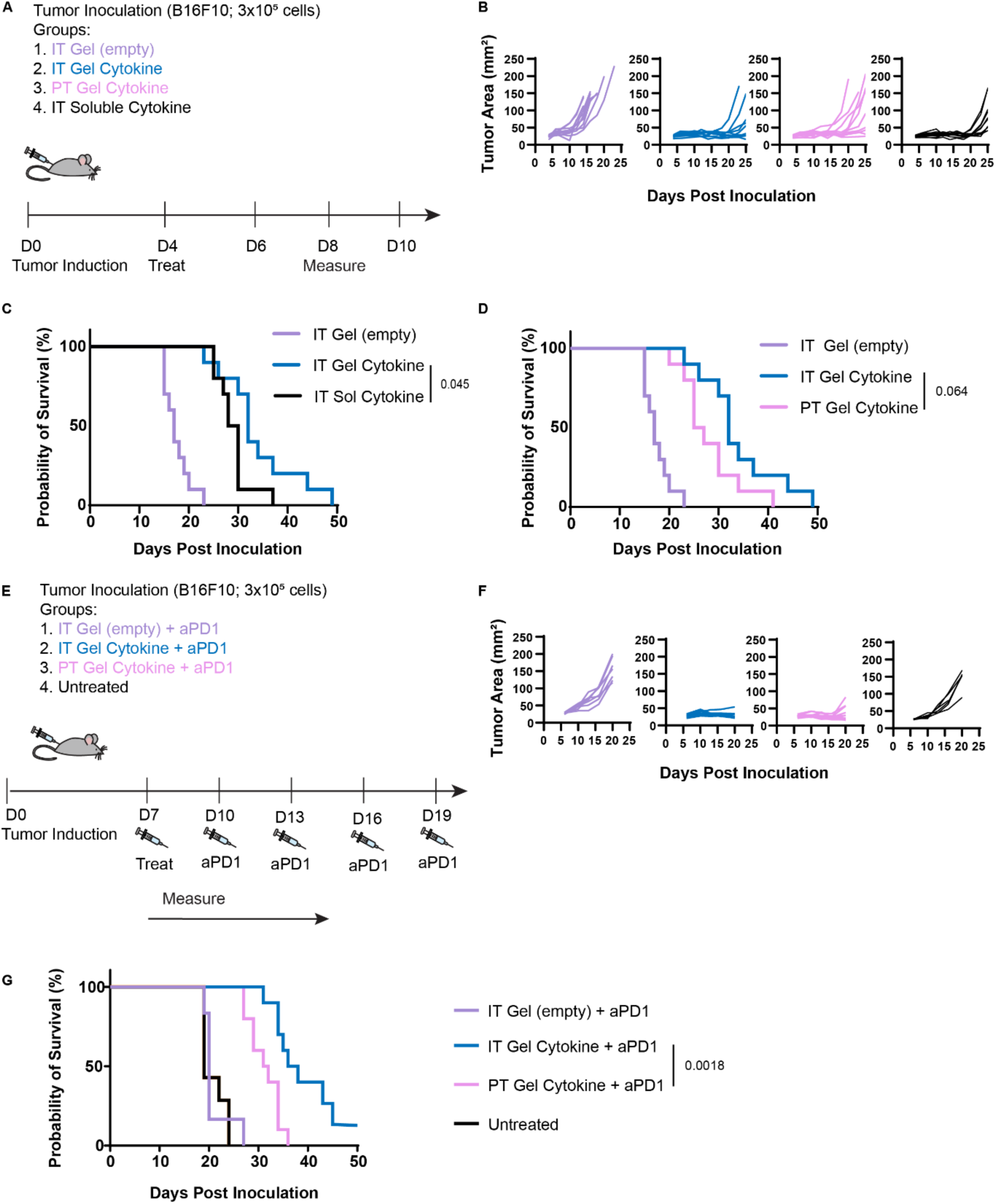
Intratumoral cytokine immunotherapy extends survival over peritumoral administration. (**A**) Experimental overview of tumor induction, treatment, and measurement timeline, N=10 for all groups. (**B**) Tumor growth over 25 days post tumor inoculation for mice treated with empty gel, intratumoral gel, peritumoral gel, or intratumoral soluble containing 5 μg of IL-12, left to right, respectively. (**C**) Survival curve comparing therapeutic effect of intratumoral gel to intratumoral soluble administration of cytokine. (**D**) Survival curve comparing therapeutic effect of intratumoral gel to peritumoral gel of cytokine. (**E**) Experimental overview of tumor induction, treatment, and measurement timeline, N=7 for vehicle control, N=6 for untreated, N=10 for all others. Cytokine treatment consisted of 5 μg IL-2 and 20 μg IL-12. (**F**) Tumor growth curves through day 20 post inoculation of intratumorally administered empty gel with ICB, intratumorally administered PNP hydrogel with cytokines in combination with ICB, untreated, and peritumorally administered PNP hydrogel with cytokines in combination with ICB, from left to right, respectively. (**G**) Probability of survival through 50 days post tumor inoculation. Survival curves were compared by log-rank Mantel-Cox test.

In this monotherapy study, we first observed that IT administration of IL-12 with PNP hydrogel led to better control over tumor growth through 25 days post tumor induction compared to soluble and PT administration of the same therapy (**Fig. 2B**). Indeed, IT PNP hydrogel administration resulted in significantly longer survival of mice compared to soluble IT administration of the same treatment, with median survival extending to 32 days from 29 days for the control treatment (*p* = 0.045, **Fig. 2C**). Further, the group receiving IT PNP hydrogel administration had survivors through day 49 post tumor induction compared to day 39 for mice receiving IT soluble cytokine. We also compared the survival benefit between mice treated with PT and IT hydrogel-based cytokine formulations. Here, there was a notable extension in the median survival by almost a week, from 26 days for mice treated with PT hydrogel-based formulations to 32 days for mice treated with IT hydrogel-based formulations (*p* = 0.0642, **Fig. 2D**).

Having demonstrated the potential for IT PNP hydrogel administration of cytokines to elicit a meaningful anti-cancer response, we next sought to examine whether a more robust response could be achieved and tolerated with higher doses and combinations of cytokines. Based on doses reported in studies using other tumor-anchoring systems, (*15–17*) we performed a dose response study, delivering either 5 µg (low dose) or 20 µg (high dose) of IL-12 on day 7 after inducing B16F10 tumors (N=8 for all groups), monitoring weight loss as an indicator of treatment-induced toxicity for two weeks post treatment (**Fig. S2**). Mice did not lose more than 5% of their body weight even when treated with high dose hydrogel-based cytokine formulations, and any weight loss was quickly recovered within one week post treatment. Moreover, we qualitatively observed that mice treated with high dose gels had more controlled tumor burden over the observation period than those mice treated with low dose gels.

We therefore chose to move forward examining the therapeutic efficacy of IT injected PNP hydrogels loaded with both 20 µg of IL-12 and 5 µg of IL-2, which are complementary cytokines that activate both the JAK and STAT pathways. In this treatment regime, all mice, except for those in the untreated group, also received immune checkpoint blockade (ICB) as 200 µg doses of aPD1 delivered intraperitoneally every three days for five total administrations (**Fig. 2E**). Mice receiving IT hydrogel-based cytokine formulations, in combination with ICB, had well controlled tumor burden through day 20 post tumor inoculation compared to those receiving other treatments (**Fig. 2F**). Mice receiving empty hydrogels and those that were untreated had tumors that started growing exponentially around 10 days after tumor inoculation, while 3 of 10 mice receiving the PT hydrogel-based cytokine treatment had tumors that started growing exponentially by day 20. The mice were monitored for long term survival, and only the mice receiving IT hydrogel-based cytokine formulations had long term survivors (defined as surviving ∼2x longer than any animal in the untreated control group, or 50 days), extending median survival from 32 to 37 days compared to those receiving PT administration of the same treatment (**Fig. 2G**, *p* = 0.0018).

### 2.3 Intratumoral hydrogel administration of cytokines polarizes the tumor microenvironment

Our initial observations of enhanced anti-cancer efficacy conferred by IT hydrogel-based cytokine formulations, but not PT hydrogel-based formulations, supported our hypothesis that the site of administration mediates critical changes to the tumor microenvironment. Specifically, these observations were consistent with the anticipated effects of the cargo delivered, IL-2 and IL-12, which are broadly pleiotropic cytokines that contribute to T cell proliferation, activation and cytotoxicity, potentially elicit counteracting immunosuppressive cues within the TME. To gain insight into the mechanism of action driving the observed survival benefit with IT PNP cytokines, we assessed the frequency of T-cell subsets in the tumor seven days following IT or PT administration of cytokine-loaded hydrogels. Interestingly, while the overall frequencies of CD8^+^ and CD4^+^ T cells among CD45^+^ cells were not different between the mice receiving IT vs PT injections, the composition of T-cell subsets were meaningfully skewed toward an anti-tumor phenotype in mice receiving IT PNP cytokines compared to those receiving PT PNP cytokines (**Fig. 3A)**. For instance, with respect to CD8^+^ T cell subsets, mice receiving IT cytokine therapy had a greater percentage of central memory, IFNγ-expressing, and naïve and stem-like memory cells (**Fig. 3A-G).** Similar trends were observed among the CD4^+^ T cells (**Fig. 3H-M).** Notably, IT PNP cytokine treatment elicited a significant increase in the fraction of CD4^+^ central memory T cells when compared to those generated by PT PNP cytokine treatment and a critical decrease in CD25^+^FOXP3^+^ regulatory T cells (**Fig. 3J,N,O).**

**Fig. 3.**
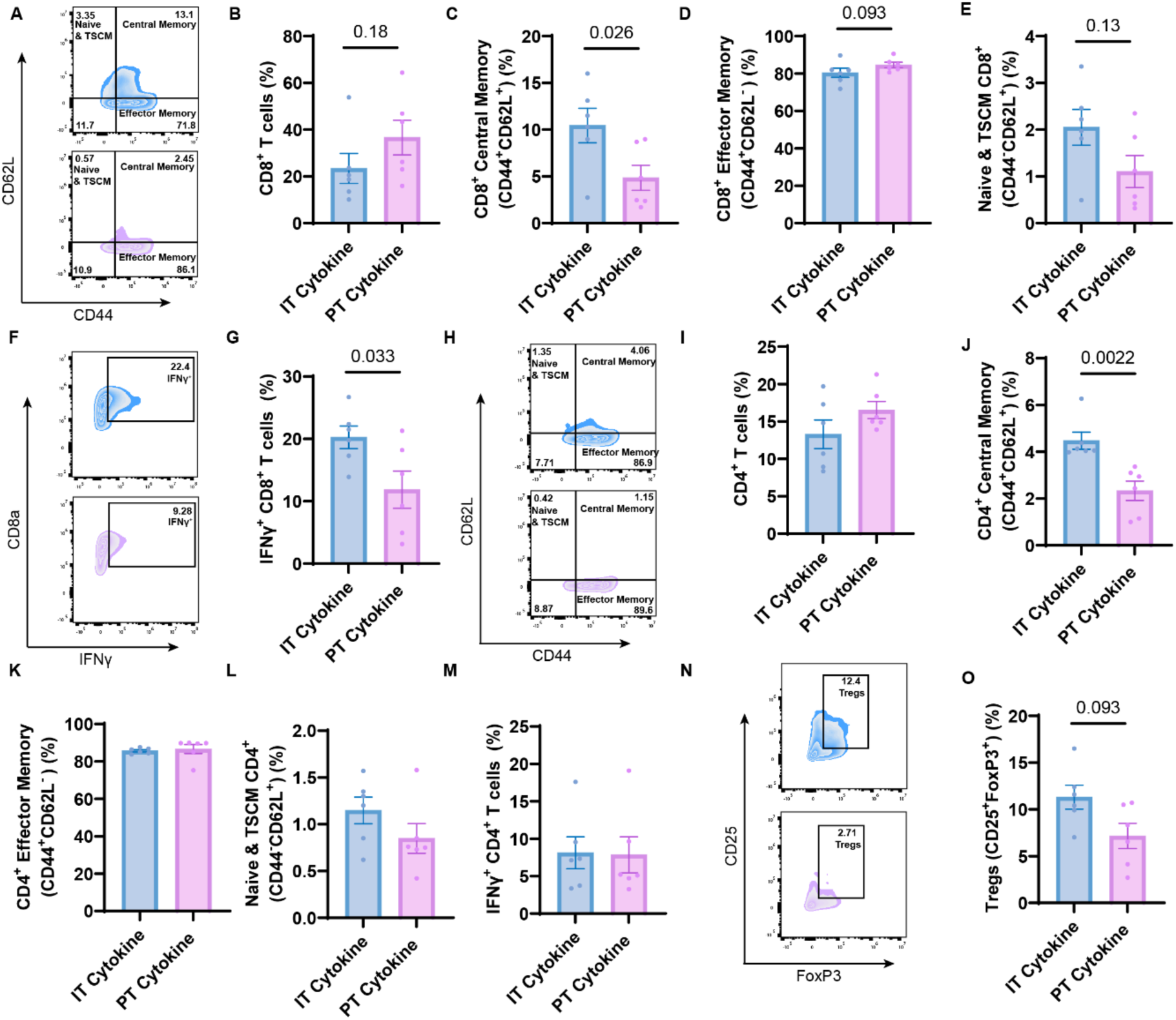
Intratumoral administration of combination cytokine immunotherapy alters the T-cell immunophenotype of the tumor. (**A**) Representative population of CD8^+^ T cells gated on CD62L and CD44 for subset phenotypes from murine B16F10 tumors; treatments included IT PNP cytokine (5 μg IL-2 + 20 μg IL-12) in blue and PT PNP cytokine (5 μg IL-2 + 20 μg IL-12) in pink. (**B**) Quantification of percentage of CD8^+^ T cells. (**C**) Quantification of percentage of CD44^+^CD62L^+^ central memory T cells. (**D**) Quantification of CD44^+^CD62L^−^ effector memory T cells. (**E**) Quantification of percentage of CD44^−^CD62L^+^ T cells. (**F**) Representative populations of interferon-gamma producing CD8^+^ T cells gated on CD8a and IFNγ. (**G**) quantification of percentages of IFNγ CD8^+^ T cells. (**H**) Representative population of CD4^+^ T cells gated on CD62L and CD44 for subset phenotypes from murine B16F10 tumors. (**I**) Quantification of percentages of CD4^+^ T cells. (**J**) Quantification of percentages of CD44^+^CD62L^+^ central memory T cells. (**K**) Quantification of percentages of CD44^+^CD62L^−^ effector memory T cells. (**L**) quantification of percentages of CD44^−^CD62L^+^ T cells. (**M**) Quantification of percentages of IFNγ^+^ CD4^+^ T cells. (**N**) Representative populations of FOXP3^+^ regulatory T cells gated on CD25 and FoxP3. (**O**) Quantification of percentages of regulatory T cells. N=6 for all groups. Tumor excised 7 days after treatment. Data reported as mean +/− SEM. Values reported as percentages of CD45^+^ cells (CD4^+^ and CD8^+^ T cells) or of parent T-cell population. Statistics determined by two-tailed Mann Whitney test in GraphPad prism.

Assessing important myeloid populations, we also observed that IT PNP cytokine treatment augmented the fraction of MHCII^+^CD11c^+^ activated antigen-presenting dendritic cells (DCs), and it particularly increased the percentage of inflammatory DCs (iDCs) when compared to PT PNP cytokines. In contrast, PT PNP cytokine treatments elicited higher fractions of cDC1s and cDC2s in the tumor (**Fig. S4A-D)**. Other populations, including B cells and natural killer (NK) cells, were similar across treatment groups in the tumors (**Fig. S4E-H**). We concomitantly examined important lymphoid and myeloid populations in the tdLN on day 7 post treatment. Here, differences in T-cell subsets were marginal, with a modest increase in the fraction of cytotoxic CD8^+^ T cells, NK cells, and MHCII^+^CD11c^+^ DCs when treated IT compared to PT (**Fig. S5A-K)**. Of note, IT PNP cytokine administration reduced the fraction of cDC2s compared to PT administration **(Fig. S5L)**.

Recognizing that the cytokine cargo has pleiotropic effects that may occur across multiple timescales, we sought to capture the immune landscape of the tumor microenvironment and the tdLNs earlier after treatment. We therefore conducted a similar analysis to assess the frequency of T-cell subsets three days post-treatment. At this early time point, we observed fewer differences in cell phenotypes between the two treatment groups, though IT PNP treatment drove an increase in the percent of cDC1s and cDC2s in the tumor while PT administration increased the representation of iDCs and NK cells (**Fig. S6A-H).** We quantified frequencies of important lymphoid and myeloid cells in the tdLNs on day three (**Fig. S7A-L)** and most notably found that IT administration of hydrogel-based cytokine formulations resulted in a slightly increased frequency of effector memory CD8^+^ T cells, a decrease in naïve and stem-like memory CD8^+^ T cells, an increase in cDC1s and a decrease in cDC2s (**Fig. S7C,D,I,K,L**).

Collectively, these results suggest that IT PNP cytokine treatments increased the antigen-specificity and proliferative capacity of T cells in the tumor microenvironment by augmenting the percentage of antigen exposed, self-renewing and migratory central memory T cells among both CD8^+^ and CD4^+^ populations. These subsets have been shown to be particularly powerful in conferring anti-tumor immunity through the production of higher levels of cytokines and longer persistence than effector memory subsets (*31–33*). These changes in T-cell subsets may be generated by shifts in the dendritic cell populations, where IT PNP cytokine treatments increased the percentage of cDC1s and cDC2s in the tdLNs at just three days post treatment, potentially priming the expansion of central memory T cells observed by day 7. Finally, IT PNP cytokine treatments resulted in an increase in IFNγ producing CD8^+^ T cells, which has direct anti-cancer consequences, while decreasing deleterious immunosuppressive regulatory T cells.

### 2.4 Peritumoral hydrogel administration of OX40a confers survival benefit over intratumoral administration

Having examined the effect of site of administration on anti-cancer immunity in the context of cytokine immunotherapies, we hypothesized that complementary trends would be observed when delivering certain antibodies like agonistic anti-OX40, which helps sustain T-cell activity and drives a long-term response through the clonal expansion of certain T-cell memory subsets. Since much of this expansion happens in the tdLNs where tumor antigens, T cells, and APCs interact, we hypothesized that PT administration of OX40a would be superior to IT administration.

To assess the therapeutic potential of OX40a, we induced B16F10 melanoma tumors in C57B/6 mice and treated with PNP hydrogels loaded with 10 µg of OX40a and 20 µg of CpG per dose on day 7. All mice, excluding those that were untreated, were also given 200 µg doses of aPD1 via intraperitoneal injection every three days for five administrations (**Fig. 4A**). Here, we observed that tumors remained well controlled in both groups receiving hydrogel-delivered therapies, while mice receiving only aPD1 or those that were untreated experienced rapid tumor growth, by two weeks post administration (**Fig. 4B**). Yet only PT administration of the hydrogel-based OX40a treatment resulted in long-term survivors, extending median survival from 26 days to 31 days when compared with IT administration (**Fig. 4C**).

**Fig. 4.**
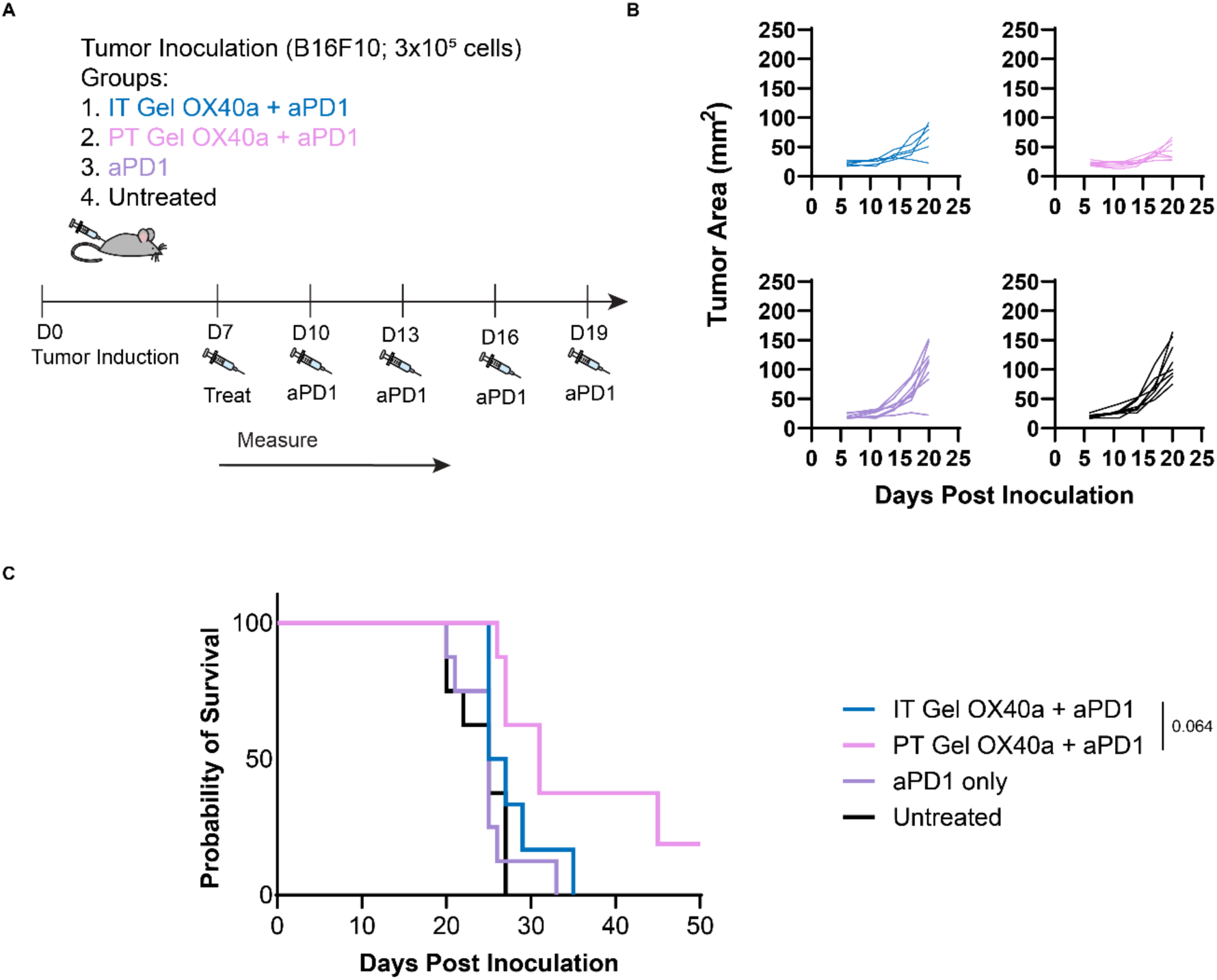
Peritumoral administration of OX40a controls tumor growth and extends murine survival in B16F10. (**A**) Experimental scheme, IT OX40a N=6, N=8 all other groups. One animal from the PT OX40a group was censored on day 31 due to the development of an ulcer. (**B**) Tumor growth curves through day 20 post inoculation of intratumorally administered OX40a with ICB, peritumorally administered OX40a with ICB, ICB only, and untreated, left to right respectively. (**C**) Probability of survival through 50 days post tumor inoculation. Survival curves were compared by log-rank Mantel-Cox test.

### 2.5 Peritumoral administration of OX40a alters tumor and lymph node immunophenotype

As with our evaluation of the impact of localization of cytokine treatments, we now sought to characterize how IT and PT administration of OX40a-loaded PNP hydrogel formulations (denoted simply IT or PT mAb) altered the immunophenotype of the tumor microenvironment and tdLNs. First, we assessed the frequency of T-cell subsets in the tumor seven days following intratumoral or peritumoral administration of OX40a-loaded hydrogels. The frequency of CD8^+^ T cells and CD8^+^ central memory T cells were similar across both treatment groups, but we found that CD8^+^ T cells were enriched for effector memory subsets when treated PT rather than IT (**Fig. 5A-D)**. We also observed that PT OX40a administration noticeably decreased the frequency of naïve and stem-like memory CD8^+^ T cells when compared to IT treatments (**Fig. 5E**), whereas both treatment groups resulted in comparable frequencies of IFNγ^+^ cytotoxic CD8^+^ T cells (**Fig. 5F, G).**

**Fig. 5.**
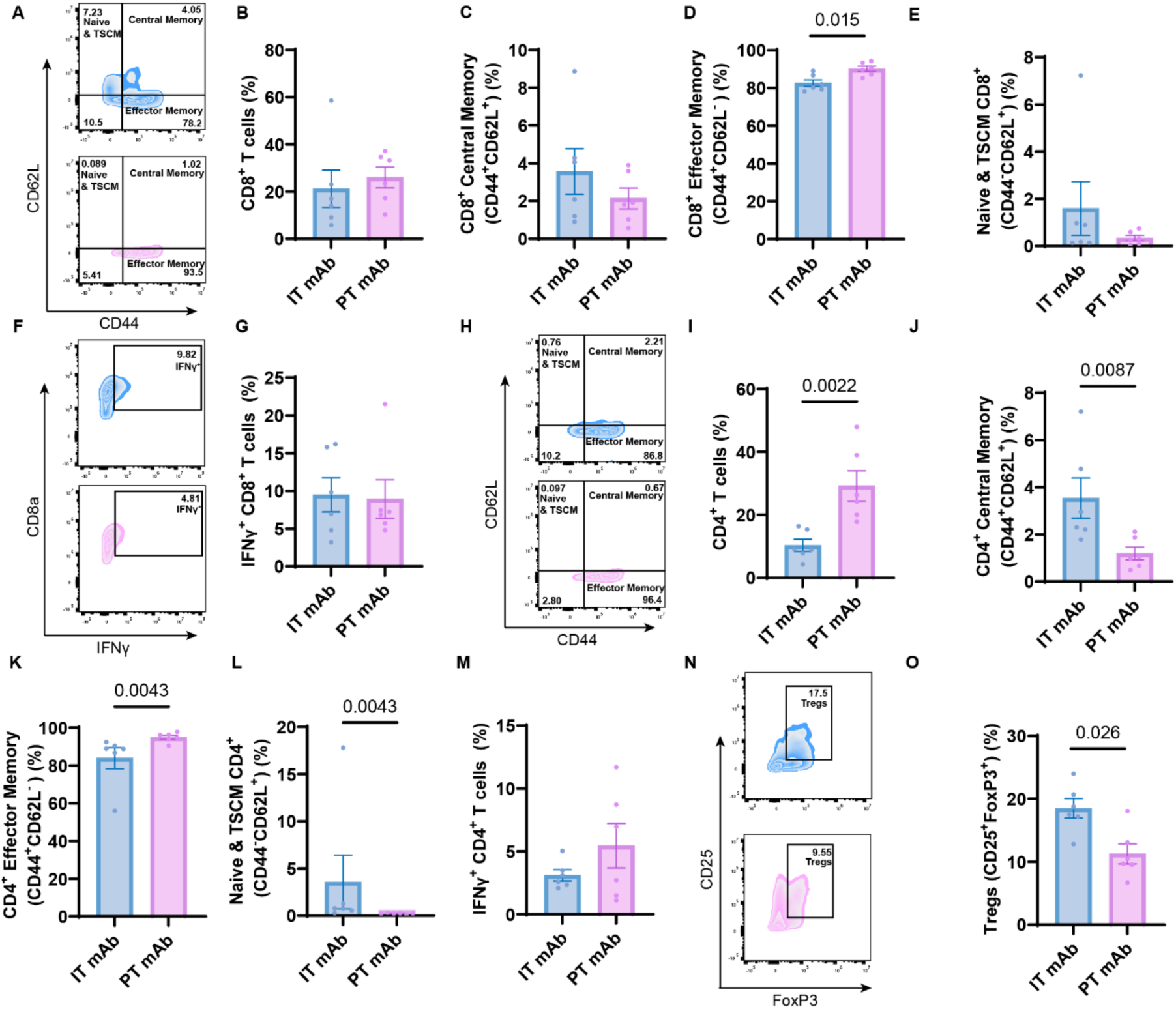
Peritumoral administration of OX40a (mAb)-hydrogels augments anti-cancer T cell phenotypes. (**A**) Representative population of CD8^+^ T cells gated on CD62L and CD44 for subset phenotypes from murine B16F10 tumors. IT PNP mAb (10 μg OX40a + 20 μg CpG) in blue and PT PNP mAb (10 μg OX40a + 20 μg CpG) in pink (**B**) Quantification of percentage of CD8^+^ T cells. (**C**) Quantification of percentage of CD44^+^CD62L^+^ central memory T cells. (**D**) quantification of CD44^+^CD62L^−^ effector memory T cells. (**E**) Quantification of percentage of CD44^−^CD62L^+^ T cells. (**F**) Representative populations of interferon-gamma producing CD8^+^ T cells gated on CD8a and IFNγ (**G**) Quantification of percentages of IFNγ^+^ CD8^+^ T cells. (**H**) Representative population of CD4^+^ T cells gated on CD62L and CD44 for subset phenotypes from murine B16F10 tumors. (**I**) Quantification of percentages of CD4^+^ T cells. (**J**) Quantification of percentages of CD44^+^CD62L^+^ central memory T cells. (**K**) Quantification of percentages of CD44^+^CD62L^−^ effector memory T cells. (**L**) Quantification of percentages of CD44^−^CD62L^+^ T cells. (**M**) Quantification of percentages of IFNγ^+^ CD4^+^ T cells. (**N**) Representative populations of FOXP3^+^ regulatory T cells gated on CD25 and FoxP3. (**O**) Quantification of percentages of regulatory T cells. N=6 for all groups. Data reported as mean +/− SEM. Values reported as percentages of CD45^+^ cells (CD4^+^ and CD8^+^ T cells) or of parent T cell population. Statistics determined by two-tailed Mann Whitney test in GraphPad prism.

Impressively, we found that PT administration of hydrogel-based OX40a formulations dramatically increased the proportion of CD4^+^ T cells and, among them, significantly boosted the fraction with an effector memory phenotype (with a corresponding reduction in the percent of central- or naïve and stem-like memory phenotypes) and those expressing IFNγ (**Fig. 5H-M**). Critically, PT administration reduced the fraction of regulatory T cells when compared to IT administration of the same therapy (**Fig. 5K**).

To complement these data, we assessed myeloid populations in the tumor on day 7 post treatment and found modest increases in frequencies of MHCII^+^ CD11c^+^ DCs, cDC2s, cDC1s, B cells, NK cells, and CD4^+^ T cells in the tumors from mice treated with PT OX40a treatments while IT treatments led to an increase in cell frequency for only iDCs and CD8^+^ T cells (**Fig. S8A-H)**.

Concurrently, we examined T-cell populations in the tdLNs on day 7 post treatment. Here, we observed similar trends, where PT administration of OX40a augmented the frequency of central memory CD8^+^ T cells and effector memory CD8^+^ T cells while decreasing the corresponding frequency of naïve- and stem-like memory CD8^+^ T cells compared to IT administration (**Fig. S9A-D**). We found a marginal increase in the frequency of helper CD4^+^ T cells but similar frequencies of CD4^+^ T cell subsets between treatment groups in the lymph nodes and similar percentages of NK cells and MHCII^+^CD11c^+^ DCs (**Fig. S9E-J**). Further, PT administration of the OX40a therapy resulted in decreased fractions of cDC1s and cDC2s in the tdLNs compared to IT administration (**Fig. S9K,L**).

Recognizing that a critical parameter of anti-cancer immunity is the generation of memory T-cell subsets and that the context in which T cells recognize antigen is paramount to T-cell expansion, contraction, and memory phases, we wanted to characterize the tumor microenvironment and tdLNs at a later stage post treatment. We again performed flow cytometric analysis on tumors and tdLNs from mice treated with OX40a-loaded hydrogels 14 days after treatment (day 21 post tumor induction). In the tumors, we noted that PT OX40a treatments increased the contribution of MHCII^+^ CD11c^+^ DCs and cDC2s (**Fig. S10A, B)** while decreasing the proportion of iDCs and cDC1s when compared to IT administration (**Fig. S10C, D**). We observed similar fractions of B cells and a small decrease in the proportion of NK cells from PT treatments compared to IT treatments (**Fig. S10E, F**). Interestingly, we found a meaningful decrease in the percentage of CD8^+^ T cells and an increase in the frequency of CD4^+^ T cells with PT OX40a treatments (**Fig. S10G, H**). Crucially, meaningful differences in frequency of lymphoid and myeloid cells persisted in the tdLNs through day 14, evidence of a sustained response (**Fig. S11A-L).**

Consistent with our findings from delivering cytokine immunotherapies, the flow cytometric analysis of the tumors and tdLNs after OX40a therapy suggests that site of administration, not just sustained locoregional delivery, is crucial to mounting a robust anti-cancer immune response. Here, PT hydrogel-based OX40a treatments elicited superior anti-tumor cellular responses compared to IT treatments, which is the exact opposite trend from cytokine immunotherapy. These results support our hypothesis that PT OX40 treatments would increase cargo exposure in the T-cell zones of lymphoid organs (*i.e.*, tdLNs) where OX40^+^ cells localize after antigen activation. In contrast to cytokine immunotherapies (which increased central memory T-cell subsets), these data suggest that the enhanced anti-tumor therapeutic efficacy of PT OX40a treatments is driven by an expansion of effector memory subsets of both CD4^+^ and CD8^+^ T cells. These effector subsets, though characterized by reduced self-renewal and longevity, acquire increased immune functionality such as rapid antigen recall and the ability to produce both inflammatory cytokines and cell lytic molecules. (*34*) Indeed, these effector subsets have been shown to correlate with positive prognosis in several cancers, including various carcinomas (*35*). Our data indicate that PT hydrogel-based OX40a treatment also resulted in dramatic expansion of CD4^+^ T cells among all T cells, decreased immunosuppressive regulatory T cells in the tumors, and prolonged T cell and APC responses. These results are consistent with the established mechanism of action of OX40 agonism, which has been shown to determine the development and longevity of CD4^+^ memory T cells by regulating clonal expansion and to regulate immunosuppression through increasing the production of cytokines, reducing regulatory T cells (*36–38*).

### 2.6 Combination of intratumorally administered cytokines and peritumorally delivered OX40a mounts robust anti-cancer response

Noting the individual efficacies of cytokines delivered IT and OX40a delivered PT, we next sought to explore possible synergy between these therapeutic approaches. We hypothesized that the combination of IT delivered cytokines and PT administered OX40a would capitalize on tumor polarization and mount a robust memory response to confer superior anti-cancer efficacy.

To assess the potency of this combination, we first induced B16F10 melanoma tumors in C57B/6 mice and treated IT with PNP hydrogels loaded with cytokines as before (5 μg IL-2 and 20 μg IL-12) with and without concomitant PT administration of PNP hydrogels loaded with 10 μg OX40a and 20 μg CpG (**Fig. 6A-C**). All groups except for the untreated control group received 250 µg of aPD1 delivered every five days for three total administrations. Reporting tumor burden through day 20 post tumor induction, we found that mice receiving hydrogel treatments had better controlled tumor growth than those that did not, consistent with our previous findings (**Fig. 6B**). Interestingly, tumor growth was better controlled in the group receiving the combination therapy of IT cytokines + PT OX40a than in the group receiving IT cytokines alone (**Fig. 6B**). This trend persisted through the duration of the study, where we found that 50% of mice treated with the IT cytokines + PT OX40a combination survived past day 50 post tumor inoculation compared, whereas no mice survived past day 41 when treated with IT cytokines and ICB alone (**Fig. 6C**). The combination therapy extended median survival almost two weeks to 46 days compared to 32.5 days for mice receiving only IT cytokine and ICB (**Fig. 6C**).

**Fig. 6.**
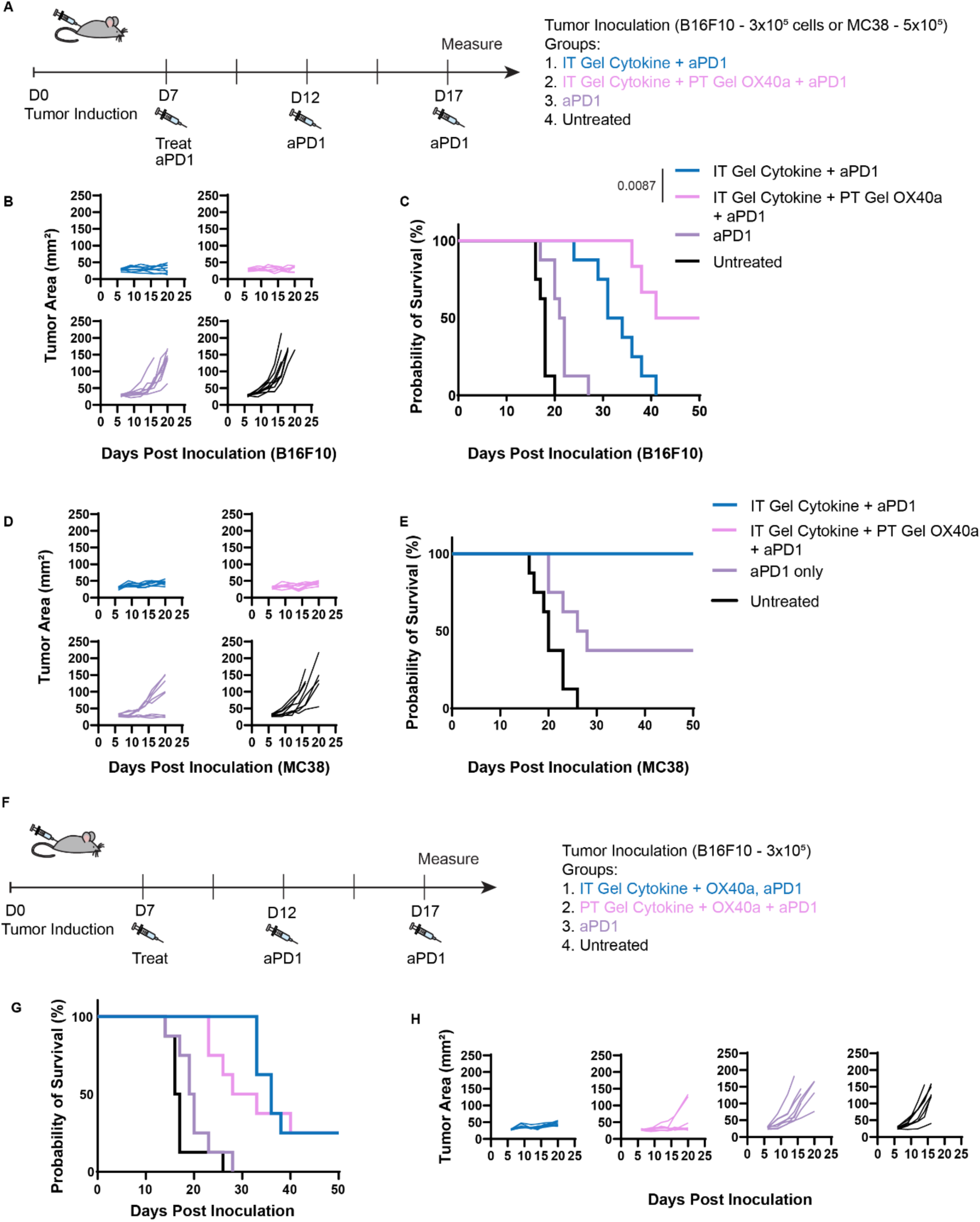
Therapeutic efficacy of intratumorally and peritumorally administered combination immunotherapy. (**A**) Experimental scheme. In the B16F10 study, N=6 for IT cytokine + PT OX40a, N=8 all other groups. In the MC38 study, N=8 for all groups. (**B**) Tumor growth curves for mice induced with B16F10 through day 20 post inoculation of intratumorally administered IL-12 and IL-2 with ICB, intratumorally administered IL-12 and IL-2 and peritumorally administered OX40a with ICB, ICB only, and untreated clockwise from top left, respectively. (**C**) Probability of survival for mice induced with B16F10 through 50 days post tumor inoculation. (**D**) Tumor growth curves for mice induced with MC38 through day 20 post inoculation of intratumorally administered IL-12 and IL-2 with ICB, intratumorally administered IL-12 and IL-2 and peritumorally administered OX40a with ICB, ICB only, and untreated clockwise from top left, respectively. (**E**) Probability of survival for mice induced with MC38 through 50 days post tumor inoculation. (**F**) Experimental scheme, N=8 for all groups (**G**) Probability of survival for mice induced with B16F10 through 50 days post tumor inoculation (**H**) Tumor growth curves for mice induced with B16F10 through day 20 post inoculation of intratumorally delivered cytokine-antibody combination, peritumorally administered cytokine-antibody combination, ICB only, and untreated, left to right respectively. Survival curves were compared by log-rank Mantel-Cox test.

To verify that these effects were tumor agnostic, we repeated the study in C57B/6 mice inoculated with 5×10^5^ MC38 murine adenocarcinoma cells and treated with the same regimens (**Fig. 6A,D,E**). Tumor burden remained well controlled over the duration of the study in both groups receiving hydrogel delivered immunotherapy, with 100% of animals being tumor free by day 50 post tumor induction in those two cohorts (**Fig. 6D, E**).

To validate that the observed synergy is a consequence of rational localization of the respective cargo, we wanted to see how efficacious the combination immunotherapy was when co-delivered at a single site. Here, we induced mice with B16F10 melanoma tumors and treated either IT or PT with a single hydrogel-based formulation comprising the entire immunotherapy combination including IL-12, IL-2, OX40a, and CpG at the same doses as before (**Fig. 6F**). Mice receiving the IT total combination therapy had a median overall survival of 36 days while those receiving the same therapy PT had a median overall survival of 30.5 days (**Fig. 6G**). Interestingly, mice receiving the IT total combination therapy had well controlled tumor burden through day 20 post tumor induction whereas 25% of those receiving the PT total combination therapy had tumors that reached exponential growth by day 23 (**Fig. 6H**). Despite the difference in immediate tumor growth, which resulted in earlier mouse euthanasia, the long-term survival of both cohorts converged by day 40 with 25% long term survivors, which is 25% fewer than when the therapies were appropriately localized in target tissues (**Fig. 6G**).

As before, we sought to characterize these responses at the cellular level. We induced B16F10 tumors, treated, and excised tumors and tdLNs for analysis by flow cytometry on day 7 post treatment. We began by assessing T-cell populations. First, we observed that the spatially distinct combination therapy comprising IT cytokines + PT OX40a resulted in the greatest frequency of CD8^+^ T cells compared to either of the monotherapies or the IT total combination. Interestingly, the IT total combination therapy resulted in a lower frequency of CD8^+^ T cells than any other treatment, indicating a deleterious effect of having all cargo delivered simultaneously into the tumor (**Fig. 7A,B**). When examining CD8^+^ T-cell subsets, we previously noted that IT cytokines drive an expansion of central memory cells, and this same trend was especially noticeable here when comparing to PT OX40a monotherapy. The central memory subsets are nevertheless recovered when the therapies are delivered together, either in a co-localized or spatially distinct fashion (**Fig. 7C**). Similarly, the IT cytokine + PT OX40a combination therapy increases the frequency of CD8^+^ effector memory cells, which was consistent with our observations for the PT OX40a monotherapy relative to IT cytokines alone, an improvement that is not realized when the therapies are co-localized (**Fig. 7D**). The expansion of these critical cell populations is reinforced by the dramatic reduction in naïve- and stem-like central memory CD8^+^ T cells in tumors from mice receiving IT cytokine + PT OX40a combination therapy, especially compared to the IT cytokine monotherapy (**Fig. 7E**). Fascinatingly, we observed that the IT cytokine + PT OX40a combination therapy elicited the greatest frequency of IFNγ producing CD8^+^ T cells compared to all other groups, salvaging the otherwise low frequency of this population observed for the PT OX40a monotherapy (**Fig. 7F,G**). These quantifications were repeated for CD4^+^ T cell subsets, where we saw that PT OX40a drove an expansion of CD4^+^ T cells relative to other treatment groups (which is mostly preserved in the combination therapy regimes and still a noticeable increase from IT cytokine therapy alone; **Fig. 7H,I**). While the combination therapies decreased CD4^+^ central memory populations, this behavior was complimented by a subtle increase in CD4^+^ effector memory fractions (**Fig. 7J,K**). Once again, these differences paralleled decreases in the naïve- and stem-like central memory CD4^+^ T cells (**Fig. 7L**). Most strikingly, however, was the synergistic effect the IT cytokine + PT OX40a combination immunotherapy had on IFNγ^+^ CD4^+^ T cells and regulatory T cells (**Fig. 7M-P**). The IT cytokine + PT OX40a combination therapy redeemed the IFNγ^+^ CD4^+^ T cell population relative to the PT OX40a monotherapy (and still meaningfully drove the population up from the IT cytokine therapy) while significantly decreasing the frequency of Tregs, a property not directly observed for either of the monotherapies alone. Importantly, this feature is unique to the IT cytokines + PT OX40a combination therapy and not the co-localized IT combination therapy.

**Fig. 7.**
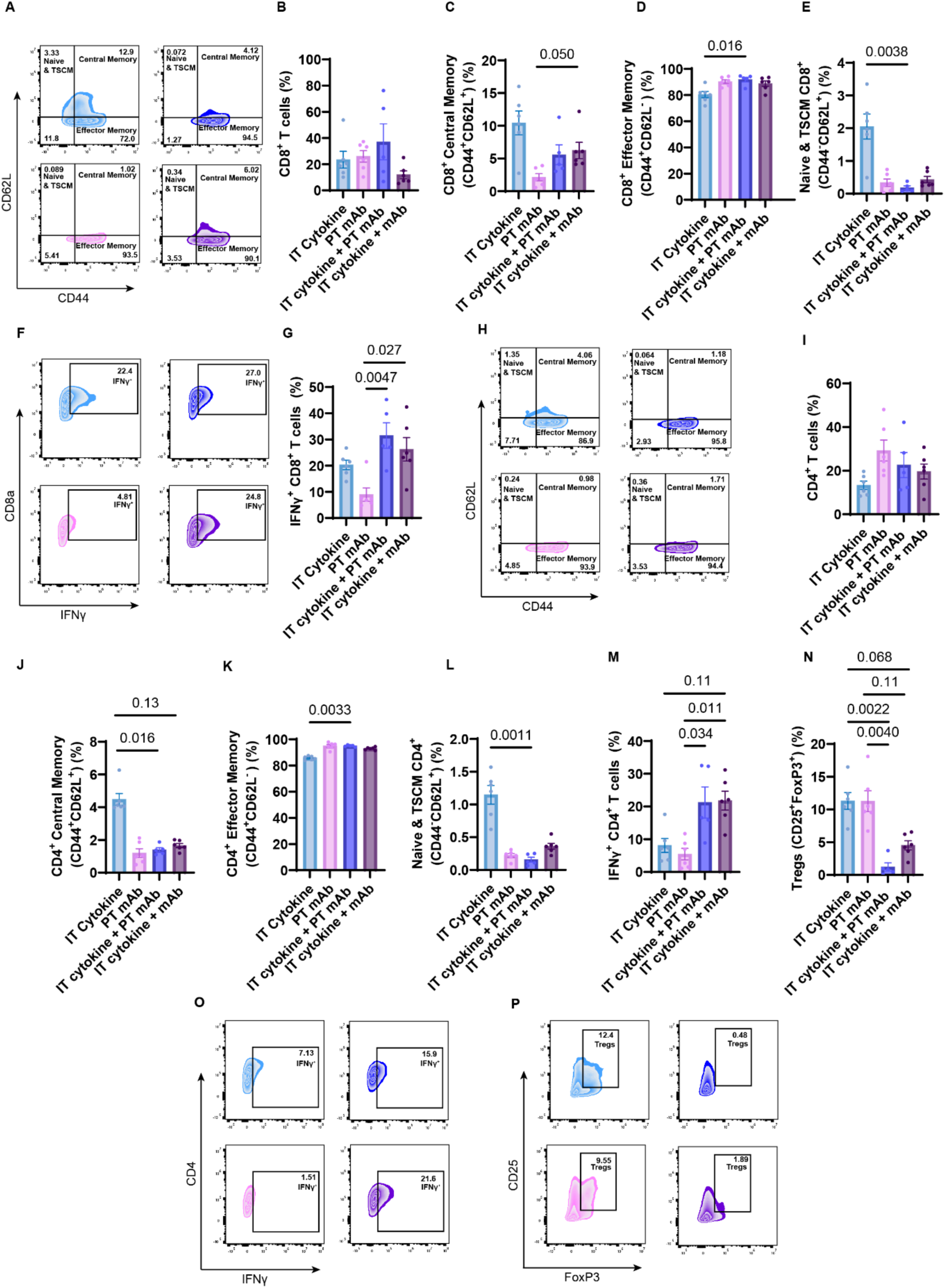
Flow cytometry of tumors day 7 after treatment with combination immunotherapies. (**A**) Representative population of CD8^+^ T cells gated on CD62L and CD44 for subset phenotypes from murine B16F10 tumors. IT PNP cytokines (5 μg IL-2 + 20 μg IL-12) in light blue, PT PNP mAb (10 μg OX40a + 20 μg CpG) in pink, IT PNP cytokines + PT PNP mAb in dark blue, IT PNP colocalized combination in purple. (**B**) Quantification of percentage of CD8^+^ T cells of CD3^+^ T cells. (**C**) Quantification of percentage of CD44^+^CD62L^+^ central memory T cells of CD8^+^ T cells. (**D**) Quantification of CD44^+^CD62L^−^ effector memory T cells of CD8^+^ T cells. (**E**) Quantification of percentage of CD44^−^CD62L^+^ naïve and stem-like T cells of CD8^+^ T cells. (**F)** Representative population of CD8^+^ T cells gated on CD8a and IFNγ for quantification of IFNγ expressing subsets from murine B16F10 tumors. (**G**) Quantification of IFNγ expressing T cells among CD8^+^ T cells. (**H**) Representative population of CD4^+^ T cells gated on CD62L and CD44 for subset phenotypes from murine B16F10 tumors. (**I**) Quantification of percentage of CD4^+^ T cells of CD3^+^ T cells. (**J**) Quantification of percentage of CD44^+^CD62L^+^ central memory T cells of CD4^+^ T cells. (**K**) Quantification of CD44^+^CD62L^−^ effector memory T cells of CD4^+^ T cells. (**L**) Quantification of percentage of CD44^−^CD62L^+^ naïve and stem-like T cells of CD4^+^ T cells. (**M**) Quantification of IFNγ expressing T cells among CD4^+^ T cells. (**N**) Quantification of CD25^+^ FoxP3^+^ regulatory T cells. (**O**) Representative population of CD4^+^ T cells gated on CD4 and IFNγ for quantification of IFNγ expressing subsets from murine B16F10 tumors (**P**) Representative population of CD4^+^ regulatory T cells gated on CD25 and FoxP3. N=6 for all groups. Values reported as percentages of CD45^+^ cells (CD4^+^ and CD8^+^ T cells) or of parent T cell population. Data reported as mean +/− SEM. Statistics determined by Kruskal-Wallis test with Dunn’s multiple comparisons test in GraphPad prism.

We continued our characterization of the tumor by examining the frequencies of various myeloid populations (**Fig. S12A-F**). Notably, the IT cytokine + PT OX40a combination therapy resulted in a moderate decrease in the proportion of MHCII^+^ CD11c^+^ DCs relative to the monotherapies, a moderate decrease in the percentage of cDC2cs relative to PT OX40a monotherapy, an increase in iDCs compared to PT OX40a monotherapy, and a decrease in cDC1s compared to both monotherapies (**Fig. S12A-D).**

Next, we characterized T-cell populations and their subsets in the tdLNs on day 7 post treatment, as before. Here, we found that the IT cytokine + PT OX40a combination therapy significantly increased the percentage of CD8^+^ T cells compared to PT OX40a monotherapy while the co-localized IT combination therapy did not (**Fig. 8A,B**). The IT cytokine + PT OX40a combination therapy resulted in a decrease in the frequency of central memory CD8^+^ T cells but dramatically increased the frequency of effector memory CD8^+^ T cells, a population which was not particularly augmented by any therapy alone (**Fig. 8A,C,D**). Both combination groups resulted in a consistent decrease in the naïve- and stem-like memory CD8^+^ T cells, though the IT cytokine + PT OX40a combination therapy did so more significantly when compared to IT cytokines alone (**Fig. 8E**). Further, in the case of the CD4^+^ T-cell population and its subsets (**Fig. 8F-J**), we noted the most impressive skew in the effector memory CD4^+^ population in the IT cytokine + PT OX40a combination therapy, which elicited significant increases frequency compared to the monotherapies.

**Fig. 8.**
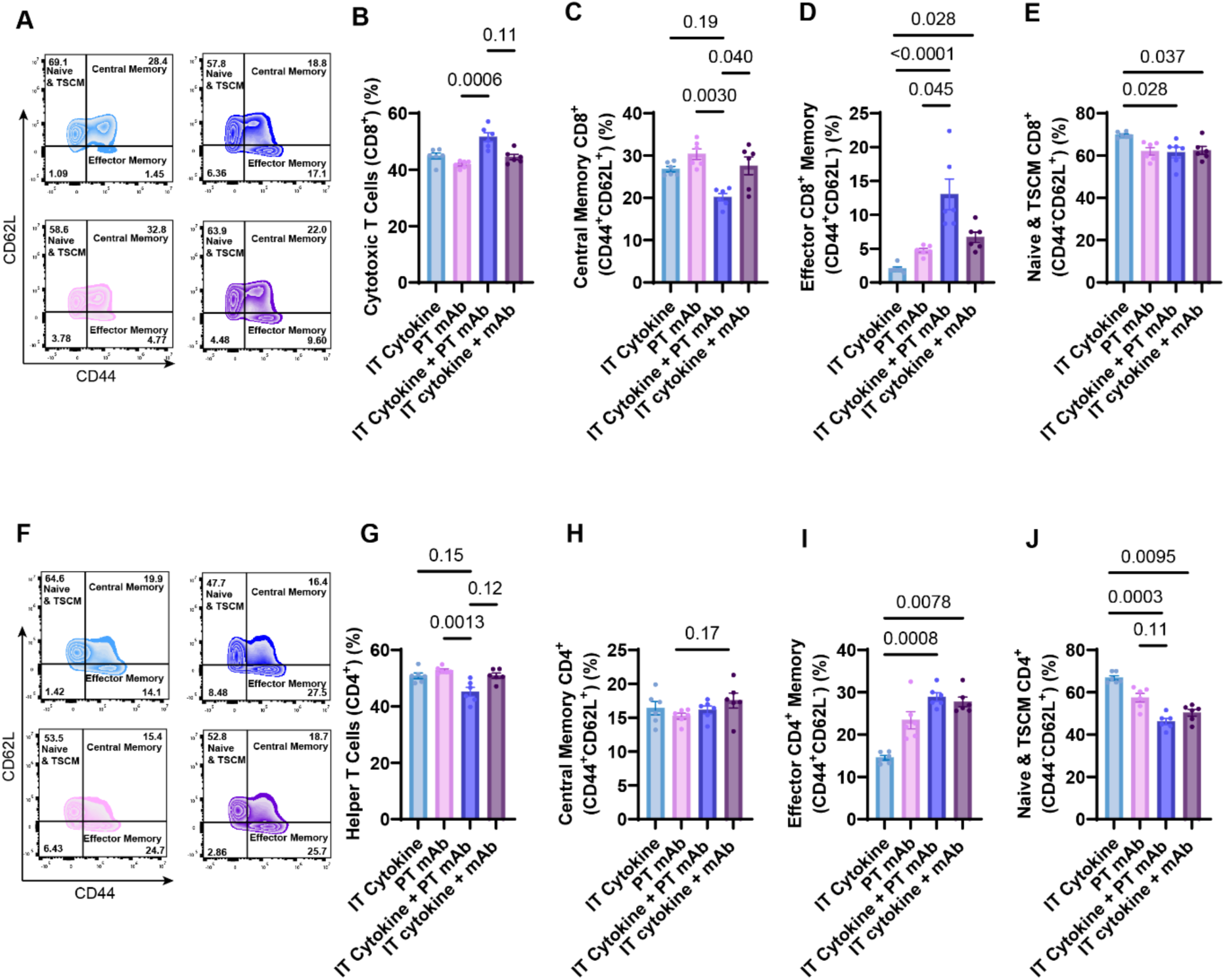
Flow cytometry of tumor draining lymph nodes day 7 after treatment with combination immunotherapies. (**A**) Representative population of CD8^+^ T cells gated on CD62L and CD44 for subset phenotypes from murine B16F10 tumors. IT PNP cytokines (5 μg IL-2 + 20 μg IL-12) in light blue, PT PNP mAb (10 μg OX40a + 20 μg CpG) in pink, IT PNP cytokines + PT PNP mAb in dark blue, IT PNP colocalized combination in purple (**B**) Quantification of percent cytotoxic T cells (CD8^+^) of CD3^+^ T cells. (**C**) Quantification of percentage of CD44^+^CD62L^+^ central memory T cells of CD8^+^ T cells (**D**) Quantification of CD44^+^CD62L^−^ effector memory T cells of CD8^+^ T cells. (**E**) Quantification of percentage of CD44^−^CD62L^+^ naïve and stem-like memory T cells of CD8^+^ T cells. (**F**) Representative population of CD4^+^ T cells gated on CD62L and CD44 for subset phenotypes from murine B16F10 tumors (**G**) Quantification percent helper T cells (CD4^+^) of CD3^+^ T cells. (**H**) Quantification of percentage of CD44^+^CD62L^+^ central memory T cells of CD4^+^ T cells (**I**) Quantification of CD44^+^CD62L^−^ effector memory T cells of CD4^+^ T cells. (**J**) Quantification of percentage of CD44^−^CD62L^+^ naïve and stem-like memory T cells of CD4^+^ T cells. N=6 for all groups. Values reported as percentages of CD45^+^ cells (CD4^+^ and CD8^+^ T cells) or of parent T cell population. Data reported as mean +/− SEM. Statistics determined by Kruskal-Wallis test with Dunn’s multiple comparisons test in GraphPad prism.

## 3. Discussion

The discovery and adoption of cancer immunotherapies, notably immune checkpoint inhibitors, has revolutionized the standard of care in cancer treatment and significantly improved outcomes for some patients. These successes notwithstanding, many approaches still fail to induce favorable response rates in a majority of patients and are associated with undesirable immune related adverse events (*39, 40*). Advancements in characterizing the cancer immunity cycle have, in turn, demonstrated the need for next generation cancer immunotherapy to leverage combination approaches to uniquely overcome multiple bio-orthogonal methods of immune evasion (*9*).

In seminal work by Moynihan et al., it was demonstrated that successful combination approaches must engage both innate and adaptive immunity to eradicate fully established tumors and mount a robust memory anti-cancer response (*22*). We have recently reported that an injectable hydrogel depot technology can redistribute exposure of delivered molecules compared to soluble systemic administration, rendering CD40a combination immunotherapy tractable by significantly reducing toxicities while improving efficacy by increasing accumulation in the tumor-draining lymph node and reducing systemic exposure (*27*). This study demonstrated that site of administration is critical to maximizing the therapeutic efficacy of different immune agonists based on their mechanism of action. Here, we demonstrated the capacity of this hydrogel depot technology to selectively deliver distinct immunomodulating cargo to the tumor and tumor draining lymph nodes to promote anti-cancer immunity.

Many reports have characterized the benefits of intratumoral anchoring of cytokines to improve efficacy while reducing toxicity. Momin *et al.* demonstrated that the combination of engineered IL-2 and IL-12 that binds to collagen can potentiate both checkpoint blockade and tumor specific antibodies to prolong survival and clear tumors in multiple syngeneic murine tumor models (*17*). Our studies corroborate these previous reports, where intratumoral administration of combination cytokine immunotherapy polarized T-cell phenotypes and increased INFy^+^ CD8^+^ T-cell populations in the tumor microenvironment, which underlies cytotoxic tumor clearance (*41*), significantly more strongly than peritumoral administration (**Fig. 3g**). This effector response may be driven specifically by the combination of IL-2 and IL-12, whereby IL-2 upregulates the IL-12 receptor and prepares effector cells for enhanced Th1 signaling, increasing production of anti-tumor cytokines like IFNγ and the proliferation of effector cells. Interestingly, intratumoral administration of these cytokines resulted in significant expansion of central memory subsets of both CD4^+^ and CD8^+^ T cells compared to peritumoral administration of the same therapy (**Fig. 3C, J**). Central memory T cells have enhanced survival and proliferative potential, providing a reservoir of antigen-exposed T cells ready to expand in response to challenges (*42*). While the exact role of different T-cell memory subsets remains an area of active investigation, several studies have suggested that central memory T cells confer superior anti-tumor immunity compared with effector memory T cells (*31, 43*). Collectively, central memory T cells are well positioned to respond to secondary antigen exposure and prolonged survival in lymphoid structures, suggesting that the enhanced contribution of central memory T cells in the intratumoral cytokine treatments we reported may confer an advantage in protecting against metastasis or relapse.

Importantly, we reported opposite trends with OX40a treatments. OX40 agonism has been shown to critically promote T effector cell activation and inhibit T regulatory cell function, encouraging especially the expansion and survival of CD4^+^ T cells, properties which have been shown crucial for eradicating local, distal, and spontaneous malignancies (*25*). In this work, we show peritumoral administration of OX40a most notably augmented T effector memory cell subsets (both CD8^+^ and CD4^+^), CD4^+^ T cells (as a fraction of all CD45^+^ cells), and downregulated immunosuppressive regulatory T cells compared to intratumoral administration of the same therapy (**Fig. 5D, I, K, O**). While CD8^+^ T cells have canonically been the focus for cancer immunotherapy due to their cytotoxic effector functions, recent reports suggest that CD4^+^ T cells play a pivotal role in directing the anti-tumor immune response and that local enrichment of their frequency correlates with survival in high risk patients (*44–46*).

The process of T cell differentiation and drivers of T cell commitment to specific cell subsets is clearly complex and regulated in both space and time. Indeed, a robust anti-cancer response appears to require a local pro-inflammatory and cytotoxic response in the tumor, characterized by high levels of IFNγ producing T cells, low levels of regulatory T cells, a pool of T cells that have acquired effector function, and a reservoir of highly proliferative T cells to maintain the longevity of the immune response. Here, we demonstrated that site-specific therapeutic administration could be rationally employed to increase exactly these desired T-cell subsets in the tumor microenvironment and tumor draining lymph node. For example, IT cytokines alone augmented the central memory T cell response and CD8^+^ T cells while PT OX40a expanded the effector memory and CD4^+^ T cell responses. Independently, neither therapeutic approach mounted a sufficiently efficacious response to result in a meaningful number of long-term survivors. Yet combination therapy of IT cytokines + PT OX40a elicited robust central and effector memory T cell responses while significantly reducing immunosuppressive regulatory T cell responses, resulting in 50% long term survivors in the highly immunosuppressive B16F10 melanoma model and 100% tumor regression in the MC38 adenocarcinoma model.

These results underscore the need to target therapy in a rational, spatiotemporal fashion. Localization to the tumor microenvironment may be insufficient to fully capitalize on a therapy’s potential depending on its mechanism of action. Central memory T cell populations traffic to the lymph node where they interact with tumor-antigen presenting dendritic cells, expand, and acquire effector function. The presence of the correct co-stimulatory cues can be deterministic to the ultimate composition of the expanded T cell population and subsequent anti-cancer immune response. We hypothesize that cytokine driven expansion of central memory T cells is complimentary to OX40 agonized lymph node interactions that drive a more robust effector memory T cell population. As suggested by the results of adoptive immunotherapy studies, a heterogeneous population of central memory and effector memory T cell populations may potentiate the best clinical outcomes (*43, 47*). Indeed, combining intratumoral cytokine-loaded hydrogels with peritumoral OX40a-loaded hydrogels resulted in synergistic effects that altered the immune landscape of the tumor microenvironment and tumor draining lymph nodes in distinct ways not captured by either monotherapy alone, and importantly, these effects were not recapitulated by co-delivery of the immune agonists at a single site. The combination therapy resulted in an entirely unique tumor immunophenotype consisting of both expanded central memory and effector memory T-cell subsets, increased IFNγ producing CD4^+^ and CD8^+^ T cells, and nearly complete regression of immunosuppressive regulatory T cells, results that realized superior efficacy compared to either of the leading monotherapies alone.

Overall, we have demonstrated a strategy for site-specific immunotherapy administration that bridges innate and adaptive immune pathways toward a wholly endogenous, antigen-agnostic anti-cancer response. This strategy of combining intratumoral and peritumoral injection depends only on the mechanism of action of the selected cargoes and not the expression of tumor or ECM specific motifs onto which the cargo will bind, thereby representing a facile and translatable way to explore many promising combination immunotherapies. In this work we limited our evaluation to one such combination immunotherapy, so future work will be necessary to generalize this strategy to other immunomodulatory cargo. As immuno-oncology continues to advance, additional targets are likely to be identified that should be examined with this approach, which we believe will enable the development of potent yet safe immunotherapy drug products. Nevertheless, this work demonstrated that our injectable hydrogel depot technology is an emerging immunoengineering system that can potentiate synergistic combination immunotherapy and address unique immunological questions regarding the precise spatiotemporal delivery of potent immune agonists.

## 4. Materials and Methods

### Study Design

The objective of this study was to elucidate whether sustained, site-specific administration of cytokine and antibody immunotherapy, enabled by an injectable hydrogel depot technology, could improve the therapeutic efficacy of these monotherapies and enable unique combination therapies. Our study design included characterization of cargo accumulation with in vivo imaging of excised tumors and tumor-draining lymph nodes from mice treated either intratumorally or peritumorally, survival monitoring, and characterization of the immune cells from tumors and tumor-draining lymph nodes of treated mice. Sample sizes for imaging were n=4, while for all other experiments sample sizes were n=6-10. In survival and flow cytometry studies, tumors were measured the day before treatment to ensure that within randomized groups initial tumor burdens were similar. Thereafter, treatments were randomized, and animals were cage blocked when the number of groups permitted. Data was collected until humanely defined endpoints specified as a threshold of tumor burden or weight loss as approved by Institutional Animal Care and Use guidelines. All animal studies were performed in accordance with the National Institutes of Health guidelines and the approval of Stanford Administrative Panel on Laboratory Animal Care (protocol APLAC-32947) and all animals were housed in the animal facility at Stanford University.

### HPMC-C_12_ Synthesis

HPMC-C_12_ was prepared according to a previously reported procedure. HPMC (1.0 g) was dissolved in anhydrous NMP (45 mL) with stirring at 80 °C for 1 h. Once cooled to room temperature, 1-dodecyl isocyanate (105 mg, 0.5 mmol) and Hunig’s base, acting as the catalyst (∼3 drops), were dissolved in 5 mL of anhydrous NMP. This solution was then added dropwise to the reaction mixture, which was stirred at room temperature for 16 h. The polymer was precipitated using acetone, redissolved in Milli-Q water (∼2 wt %), and dialyzed (3 kDa MWCO) against water for 4 days. The polymer was lyophilized and then reconstituted into a 60 mg/mL solution in sterile PBS 1X.

### Synthesis of PEG-*b-*PLA

PEG-*b*-PLA was prepared as previously reported. Prior to use, commercial lactide was recrystallized in ethyl acetate, and dichloromethane (DCM) was dried via cryodistillation. Under an inert atmosphere (N_2_), PEG-methyl ether (5 kDa, 0.25 g, 50 μmol) and DBU (15 μL, 0.1 mmol) were dissolved in 1 mL of anhydrous DCM. Lactide (1.0 g, 6.9 mmol) was dissolved under N_2_ in 3 mL of anhydrous DCM. The lactide solution was then quickly added to the PEG/DBU mixture and was allowed to polymerize for 8 min at room temperature. We then quenched the reaction with an aqueous acetic acid aqueous solution. Polymer was precipitated into a 1:1 mixture of ethyl ether and hexanes, collected by centrifugation, and dried under vacuum. NMR spectroscopic data, *M*_n_, and dispersity agreed with those previously described.

### PEG-b-PLA nanoparticle formation

PEG-*b*-PLA NPs were prepared as previously described. A 1 mL solution of PEG-*b*-PLA and N_3_-PEG-*b*-PLA in 75:25 ACN:DMSO (50 mg/mL) was added dropwise to 10 mL of Milli-Q water with stirring at 600 rpm. The particle solution was purified in centrifugal filters (Amicon Ultra, MWCO 10 kDa) at 4500 RCF for 1 h and resuspended in PBS 1X to reach a final concentration of 200 mg/mL.

### PNP hydrogel formulation

HPMC-C12 and PEG-PLA were prepared as described in the supplemental methods. To prepare PNP hydrogels, HPMC-C_12_ was dissolved at 6 wt% in PBS and loaded into a 1 mL luer-lock syringe. A 20 wt% solution of PEG-PLA NPs in PBS was added to a solution of PBS with adjuvant, cytokine, and/or antibodies, depending on formulation, and loaded into a second 1 mL syringe. The two syringes were connected with a female-female luer lock elbow, with care to avoid air at the interface of the HPMC-C_12_ and nanoparticle solution, and gently mixed until a homogenous PNP hydrogel was formed. Hydrogels were formulated with final concentrations of 1 wt% HPMC-C_12_ and 5 wt% NPs. Each dose consisted of 20 µL hydrogel injection either intratumorally or peritumorally through a 27-gauge needle, as indicated. Intratumoral soluble/bolus controls were administered as a 20 µL injection in PBS through a 27-guage needle, while systemic treatments of anti-PD1 were delivered as 100 µL soluble intraperitoneal injections.

### Immunotherapy formulations

Immunotherapy injections were formulated with varying concentrations of cytokines, antibodies and adjuvants in either soluble form (in phosphate buffered saline, PBS) or in Polymer Nanoparticle hydrogels (PNP hydrogels). IL-12 was dosed at 5 or 20 μg per injection, as indicated (Sino Biological CT022-M08H). IL-2 was dosed at 5 μg per injection (Sino Biological, 51061-MNAE). Anti-OX40 antibody was dosed at 10 μg per injection (BioX Cell, BE0031). ODN 2395 (CpG) was dosed at 20 μg per injection (Invivogen, tlrl-2395). Anti-PD1 antibody was dosed at 200 or 250 μg per injection, as indicated (BioX Cell, Clone RMP1-14, BE0146).

### Mammalian cell culture

Murine B16F10 melanoma was purchased from ATCC (CRL-6475). Aliquots of MC38 murine colon adenocarcinoma were provided as a gift from the Davis Lab at Stanford and subsequently purchased from Sigma Aldrich (SCC172). Cells were maintained in RPMI media (Cytiva, SH30027.FS), supplemented with 10% fetal bovine serum (R&D Systems, S11550) and 1% penicillin-streptomycin (Themo Fisher, 15140122) prior to tumor inoculation.

### Murine tumor induction and monitoring

All animal studies were performed in accordance with the National Institutes of Health (NIH) guidelines, with the approval of the Stanford Administrative Panel on Laboratory Animal Care (protocol APLAC-32947). Seven to eight week old female C57BL/6 were purchased from Charles River and housed in the animal facility at Stanford University. Mice were allowed to acclimate in the Stanford facilities for 1 week prior to beginning experimental procedures. Right-side flanks of mice were shaven prior to injections.

MC38 flank xenografts were generated on 9-week-old female C57BL/6 mice by injecting a 50 µL of MC38 cells (10 × 10^6^ cells/mL) encapsulated in a 1 wt% alginate hydrogel subcutaneously above the right hind leg. Cell loaded alginate hydrogels were prepared using a dual syringe mixing technique as described previously. A stock solution of sterile alginate (Pronova UP LVG) was dissolved at 5wt% by adding saline to the polymers and allowing them to dissolve over 1 day at 4C. A stock solution of calcium sulfate (100 mM) was prepared in water in a large container in the form of a slurry, mixed vigorously. Calcium sulfate and cells were combined at the volume needed to reach the desired final concentration and loaded into a 1 mL syringe, while alginate (at the volume needed to reach the desired final concentration) was loaded into a separate 1 mL syringe. The two syringes were connected with a female-female luer lock elbow with care to avoid air at the polymer-solution interface, and the solutions were mixed gently until a homogenous alginate hydrogel (1 wt% alginate, 10 mM calcium sulfate) was formed.

B16F10 melanoma models were generated similarly, whereby 9-week-old female C57BL/6 mice were injected with a 50 µL alginate hydrogel loaded with B16F10 cells (6 × 10^6^ cells/mL) subcutaneously above the right hind leg.

Following tumor inoculation, mice were monitored for palpable tumors. Treatments began on day 7 after tumor inoculation, at which point tumors grew to approximately 5 mm in diameter. Treatment groups were formed to maintain consistent tumor burden at the time of treatment, resulting in groups of N = 6 – 10, depending on the study design. Mice were cage blocked or randomized depending on the study. Tumor burden was measured on D-1 before treatment to ensure fair distribution of tumors across treatment groups. Mouse weight was tracked to assess acute toxicity, with euthanasia criteria for sustained losses of more than 20% of body mass relative to the start of treatment. Tumor burden was tracked three times a week using caliper (Mitutoyo) measurements to calculate total tumor area (*L* × *W*); mice were euthanized once total tumor burden grew to 150 mm^2^ or larger.

### AlexaFluor 647 conjugation of anti-OX40 antibody

Anti-OX40 antibody was conjugated with AlexaFluor 647 NHS-ester (ThermoFisher, A20006) according to the manufacturer’s specifications. Briefly, AlexaFluor 647 NHS-ester was prepared at 10 mg/mL in anhydrous DMSO and combined with the antibody in 8 molar excess. The reaction was left to stir and incubate for 1 hour at room temperature before purification of the conjugated antibody via repeated washes through Amicon ultra centrifugal filters, 10 kDA MWCO (Millipore, UFC501008).

### *In vivo* biodistribution of anti-OX40 antibody

C57BL/6 mice were injected either peritumorally or intratumorally with 1:5 PNP hydrogels loaded with 10 µg of AF647 labeled anti-OX40 antibody. At the indicated time points, mice were euthanized with CO_2_ and either the lymph node or tumors were excised and imaged using an in vivo imaging system (IVIS Lago). Excised tissues of interest were imaged at 640/670 excitation/emission with medium binning, fstop = 8 for 2 second exposure.

### Sample preparation for flow cytometry of lymph nodes and tumors

Tumor draining inguinal lymph nodes were harvested from mice at days 3, 7, and 14 post intratumoral or peritumoral immunotherapy administration, as indicated. LNs were mechanically disrupted to single cell suspensions using Corning single-frosted micro slides (2948-75X25), passed through 70 micron filters (Celltreat, 229484), spun at 500 rcf for 5 minutes, resuspended in PBS, and counted using acridine orange/propidium iodide cell viability stain (Vitascientific, LGBD10012) and a Luna-FL dual Fluorescence cell counter (Logos biosystems). 1 million live cells per sample were transferred to a 96-well conical bottom plate (Thermo Scientific, 249570) and stained.

Tumors were similarly excised, weighed and mechanically disrupted via surgical scissors. Bulk homogenate was enzymatically digested with an enzyme mixture consisting of collagenase/hyaluronidase (STEMCELL Technologies, # 07912) and DNAse 1 (STEMCELL Technologies, #100-0762) at 0.1x:0.15x ratios, respectively, in RPMI media (Cytiva, SH30027.FS). Samples and enzymatic mixture were transferred to 5.0 mL protein lobind tubes (Eppendorf, #0030108302) and left to digest at 37C while shaking at 300 rpm for 25 minutes. Digestion was stopped by adding complete RPMI media, previously described, supplemented with 1% 0.5 M EDTA (BioVision, 2103-100). Samples were spun at 300 rcf for 10 min with brake on low, resuspended in PBS, counted and plated as described for LNs

### Staining protocol and flow cytometry

LN and tumor samples were first stained with 200uL Ghost Dye Violet 510 live/dead (Tonbo Biosciences, 13-0870-T100) for 5 minutes on ice, quenched with 100 µL FACS buffer (PBS, 3% heat inactivated FBS, 1 mM EDTA), and spun at 935xg for 2 minutes (with brake on low for tumor samples). Samples were then incubated with 50 µL anti-mouse CD16/CD32 (1:50 dilution, BD, 553142) for 5 minutes on ice before staining with 50 µL surface antibody stain for 30 min on ice. Samples were spun as before. For intracellular cytokine staining, samples were incubated in 200 µL 1x IC Fixation Buffer for 30 minutes at room temperature in the dark. Cells were spun at 500xg for 5 minutes, washed in 200 µL of 1x Permeabilization Buffer, and spun as before. Samples were then incubated in 100uL of full intracellular antibody stain diluted in 1x Permeabilization buffer for 30 minutes at room temperature in the dark. Cells were quenched in 200 µL of 1x Permeabilization buffer, centrifuged at 500xg for 5 minutes, then washed and spun once more. Samples were resuspended in 200uL FACS buffer and run on the Agilent NovoCyte Penteon Flow Cytometer in the Stanford Shares FACS Facility. Data was analyzed in FlowJo. FoxP3/Transcription Factor Staining Buffer Set (Thermo Fisher Scientific, 00-5523-00) were used for fixation and permeabilization according to the manufacturer’s instructions.

LN full antibody stain included anti-mouse CD44 (1:80 dilution; BV750; BioLegend, 103079), anti-mouse/rat XCR1 (1:160 dilution; AF647; BioLegend, 148213), anti-mouse CD11c (1:200 dilution; PE; BioLegend, 117308), anti-mouse CD19 (1:200 dilution; PE-Cy7; BioLegend, 115520), anti-mouse CD161 (NK1.1) (1:100 dilution; BV605; BioLegend, 108740), anti-mouse I-A/I-E (MHCII) (1:400 dilution; FITC; BioLegend, 107606), anti-mouse CD45 (1:400 dilution; AF700; BioLegend, 103128), anti-mouse CD8a (1:200 dilution; BV785; BioLegend, 100750), anti-mouse CD3 (1:200 dilution; PerCP/eFluor710; Invitrogen, 46003283), anti-mouse CD11b (1:100 dilution; BUV395; BD, 563553), anti-mouse F4/80 (1:125 dilution; BV421; BioLegend, 123132), anti-mouse CD4 (1:200 dilution; BUV805; BD, 612900), and anti-mouse CD62L (1:80 dilution; BUV737; BD, 612833).

Tumor surface antibody stain included anti-mouse CD25 (1:50 dilution; FITC; BioLegend, 101908), anti-mouse/rat XCR1 (1:100 dilution; AF647; BioLegend, 148213), anti-mouse CD206 (1:50 dilution; BV650; BioLegend, 141723), anti-mouse Ly-6C (1:100 dilution; BV570; BioLegend, 128030), anti-mouse Ly-6G (1:100 dilution; BV711; BioLegend, 127643), anti-mouse CD11c (1:100 dilution; PE; BioLegend, 117308), anti-mouse CD19 (1:100 dilution; PE-Cy7; BioLegend, 115520), anti-mouse CD161 (NK1.1) (1:50 dilution; BV605; BioLegend, 108740), anti-mouse I-A/I-E (MHCII) (1:200 dilution; FITC; BioLegend, 107606), anti-mouse CD45 (1:200 dilution; AF700; BioLegend, 103128), anti-mouse CD8a (1:100 dilution; BV785; BioLegend, 100750), anti-mouse CD3 (1:100 dilution; PerCP/eFluor710; Invitrogen, 46003283), anti-mouse CD11b (1:100 dilution; BUV395; BD, 563553), anti-mouse F4/80 (1:80 dilution; BV421; BioLegend, 123132), and anti-mouse CD4 (1:100 dilution; BUV805; BD, 612900). Tumor intracellular antibody stain included anti-mouse IFN-γ (1:20 dilution; APC; BioLegend, 505810) and anti-mouse FoxP3 (1:20 dilution; PE; BioLegend, 126403).

### Statistical analysis

For imaging analysis, n=4 samples were imaged per timepoint per group, and data are presented as mean +/− standard error of mean (SEM) as specified in the corresponding Fig. captions. Statistics are ordinary one-way ANOVA run in GraphPad Prism with Tukey’s multiple comparisons test.

For survival studies and flow cytometry, a sample size of n=6-10 was used and data are presented as mean +/− standard error of mean (SEM) as specified in the corresponding Fig. captions. Comparisons between two treatment groups (i.e intratumoral to peritumoral) were determined by two-tailed Mann Whitney test in GraphPad prism. Survival curves were compared by log-rank Mantel-Cox test in GraphPad prism. Comparisons between multiple groups were determined by Kruskal-Wallis test with Dunn’s multiple comparisons test in GraphPad prism. p values less than 0.2 are shown.

## 6. Acknowledgements

## Funding

This work was supported financially in part by the Goldman Sachs Foundation (administered by the Stanford Cancer Institute, SPO# 162509). JHK is thankful for the generous support of an Agilent Fellowship. A.N. is grateful for funding from the Paul and Mildred Berg Fellowship. E.L.M. was supported by the NIH Biotechnology Training Program (T32-GM008412). B.S.O. was supported by Eastman Kodak Fellowship. J.B. is thankful for a Marie-Curie fellowship from the European Union (H2020; No. 101030481). Flow cytometry was performed on an instrument in the Stanford Shared FACS Facility obtained using NIH S10 Shared Instrument Grant (1S10OD026831-01).

## Author contributions

Conceptualization: JHK, EAA

Methodology: JHK, AN, ELM, BSO, JB, EAA

Investigation: JHK, AN, ELM, BSO, JB

Visualization: JHK, AN, EAA

Supervision: JHK, EAA

Funding Acquisition: EAA

Project Administration: JHK, EAA

Writing – original draft: JHK

Writing – review & editing: JHK, AN, ELM, BSO, JB, EAA

## Competing interests

E.A.A., J.H.K., E.L.M and A.N. are listed as inventors on a pending patent application describing the technology reported in this manuscript. E.A.A. is a co-founder, equity holder, and advisor to Appel Sauce Studios LLC, which holds a global exclusive license to this technology from Stanford University. All other authors declare no competing financial interests.

## Data and materials availability

All data supporting the results in this study are available within the article and its Supplementary Information. The broad range of raw datasets acquired and analyzed (or any subsets thereof), which would require contextual metadata for reuse, are available from the corresponding author upon reasonable request.

