## Supplemental Information for "Spatially Tuned Localization of Interleukins and OX40 Agonists Enhances Synergistic Anti-Tumor Immunity"

A.

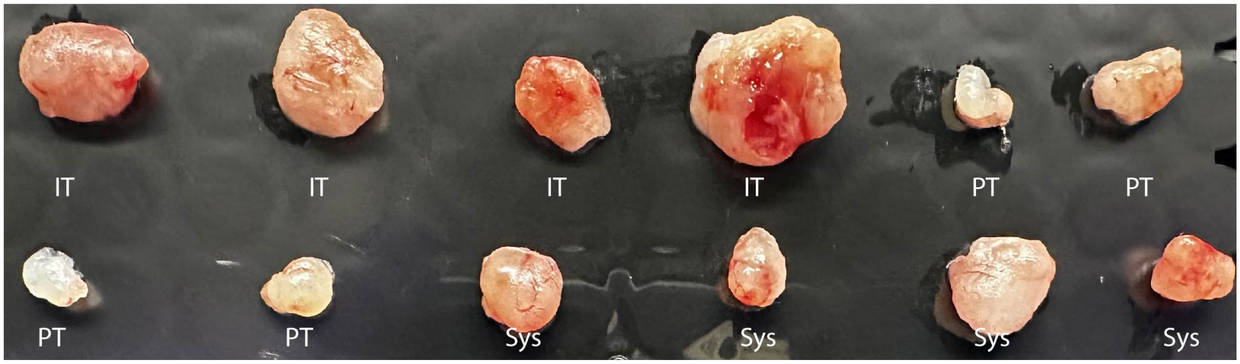


**Figure S1. Qualitative inspection of MC38 tumor burden.** Tumors were excised on day 3 after treatment for fluorescent imaging by IVIS and inspected qualitatively.

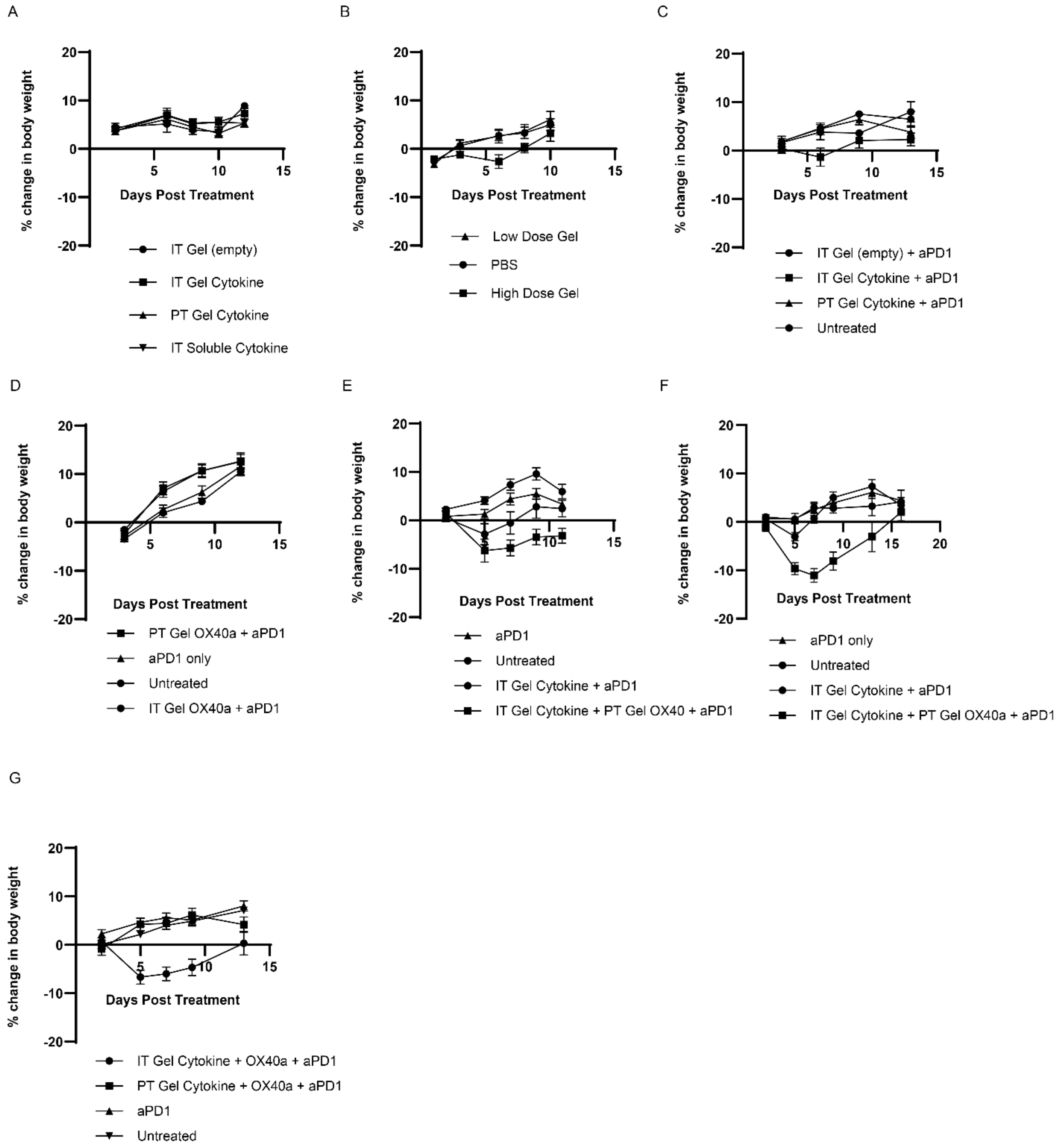


**Figure S2. Animal weight monitoring post treatment for all studies as a proxy for toxicity.** Weight loss of animals in studies corresponding to **A.** IL-12 monotherapy study reported in Figure 2A **B.** dose response study of IL-12 at 20 ug (high) and 5 ug (low) **C.** IL-12/IL-2 cytokine study reported in Figure 2E **D.** OX40a monotherapy study reported in Figure 4A **E.** Cytokine and antibody combination therapy study done in B16F10 reported in Figure 6A **F.** Cytokine and antibody combination study done in MC38 reported in Figure 6A **G.** Single-site cytokine and antibody combination study reported in Figure 6F.

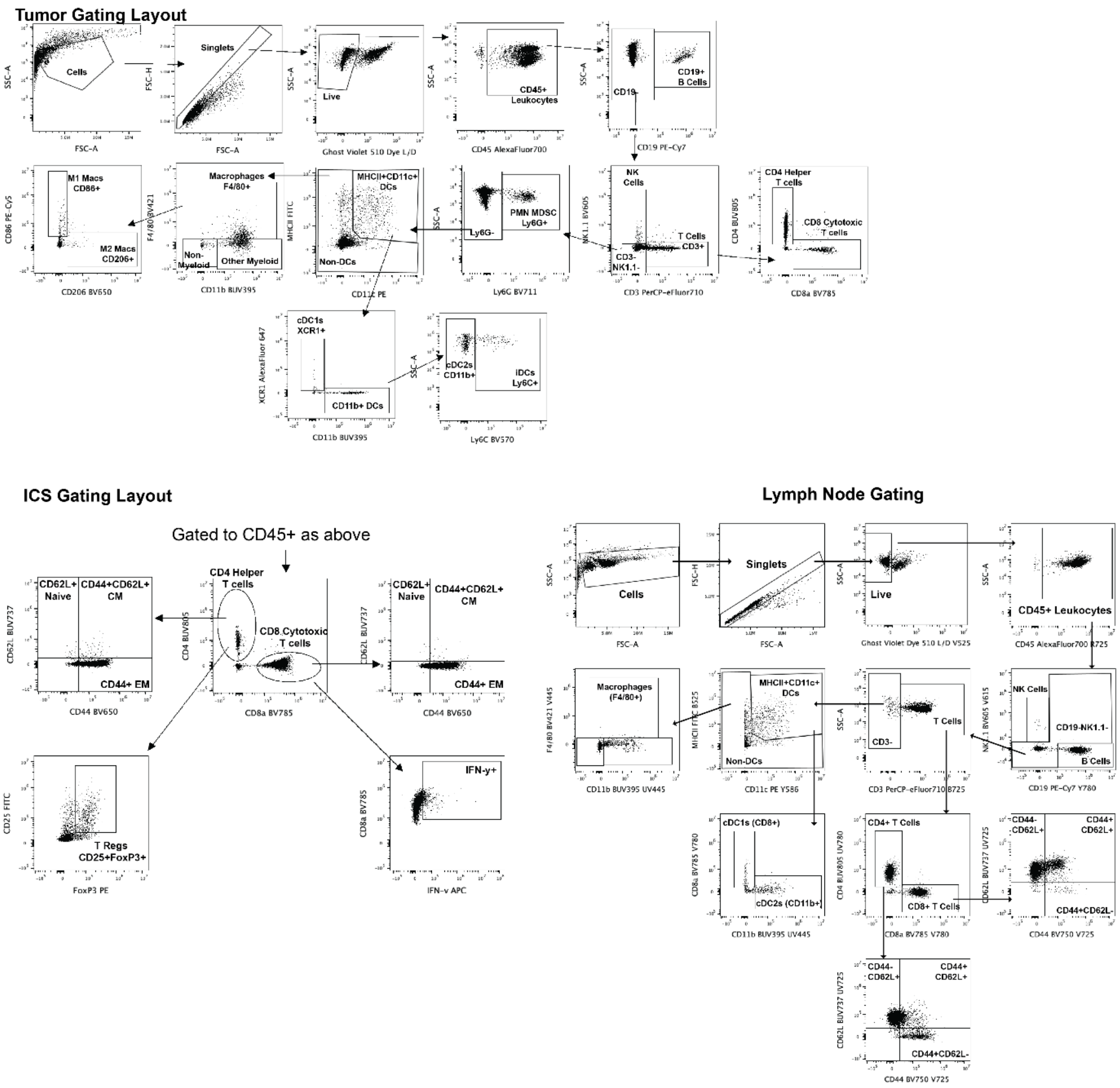


**Figure S3. Representative gating strategies for tumors, lymph nodes and intracellular (ICS) staining of tumors, clockwise respectively.**

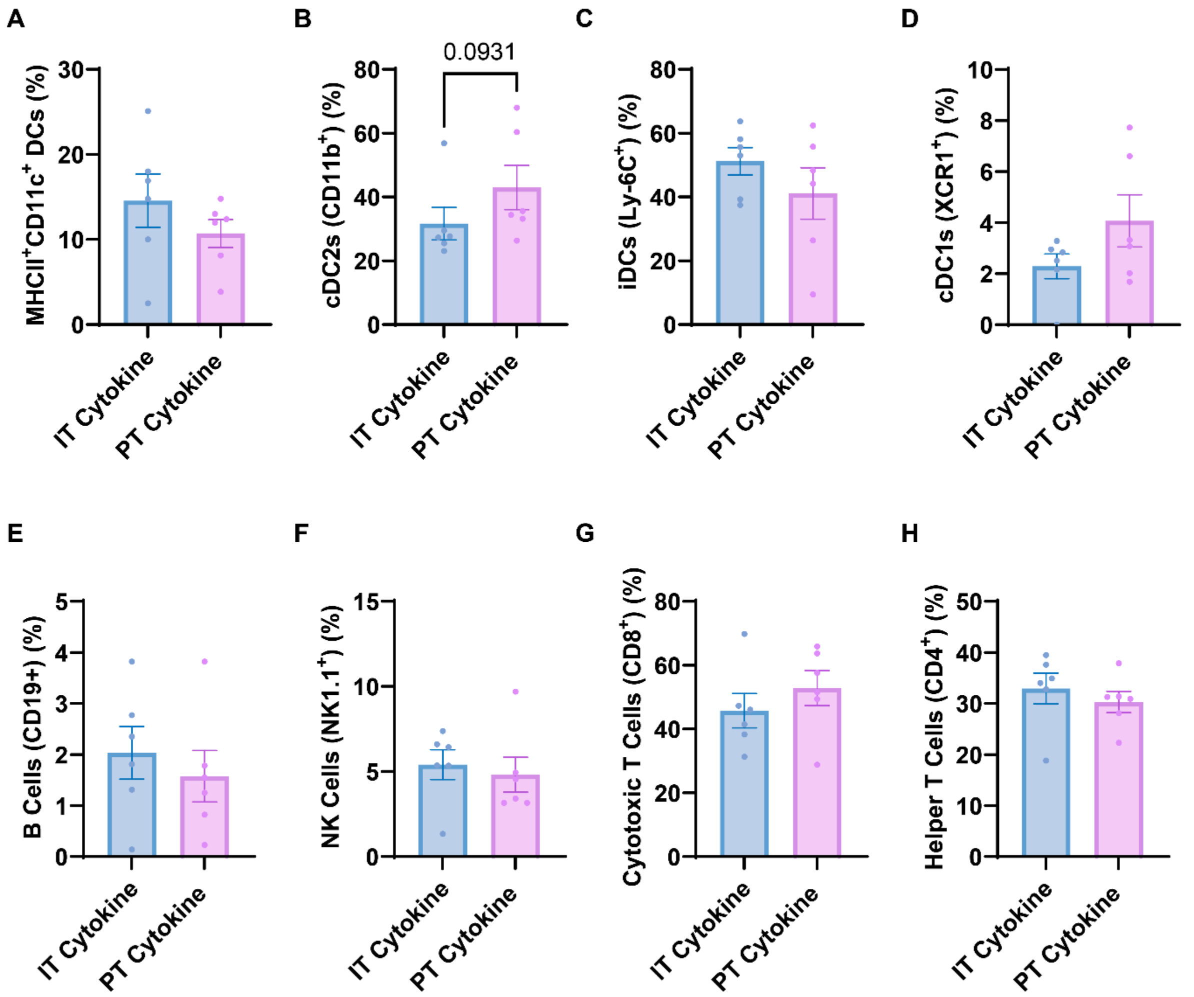


**Figure S4. Additional flow cytometry of tumors on day 7 post cytokine therapies**. **A.** Percent of MCHII$^{+}$ CD11c$^{+}$ DCs of CD45$^{+}$ cells **B.** Percent of cDC2s (CD11b$^{+}$) of MCHII$^{+}$ CD11c$^{+}$ DCs **C.** Percent of iDCs (Ly-6C$^{+}$) cells of MCHII$^{+}$ CD11c$^{+}$ DCs **D.** Percent cDC1s (XCR1$^{+}$) cells of MCHII$^{+}$ CD11c$^{+}$ DCs **E.** Percent of B cells (CD19$^{+}$) of CD45$^{+}$ cells **F.** NK cells (NK1.1$^{+}$) cells of CD45$^{+}$ cells **G.** Percent of CD8$^{+}$ T cells of CD3$^{+}$ T cells **H.** Percent of CD4$^{+}$ T cells of CD3$^{+}$ T cells. N=6 for all groups. Data reported as mean +/- SEM. Statistics determined by two-tailed Mann Whitney test in GraphPad prism.

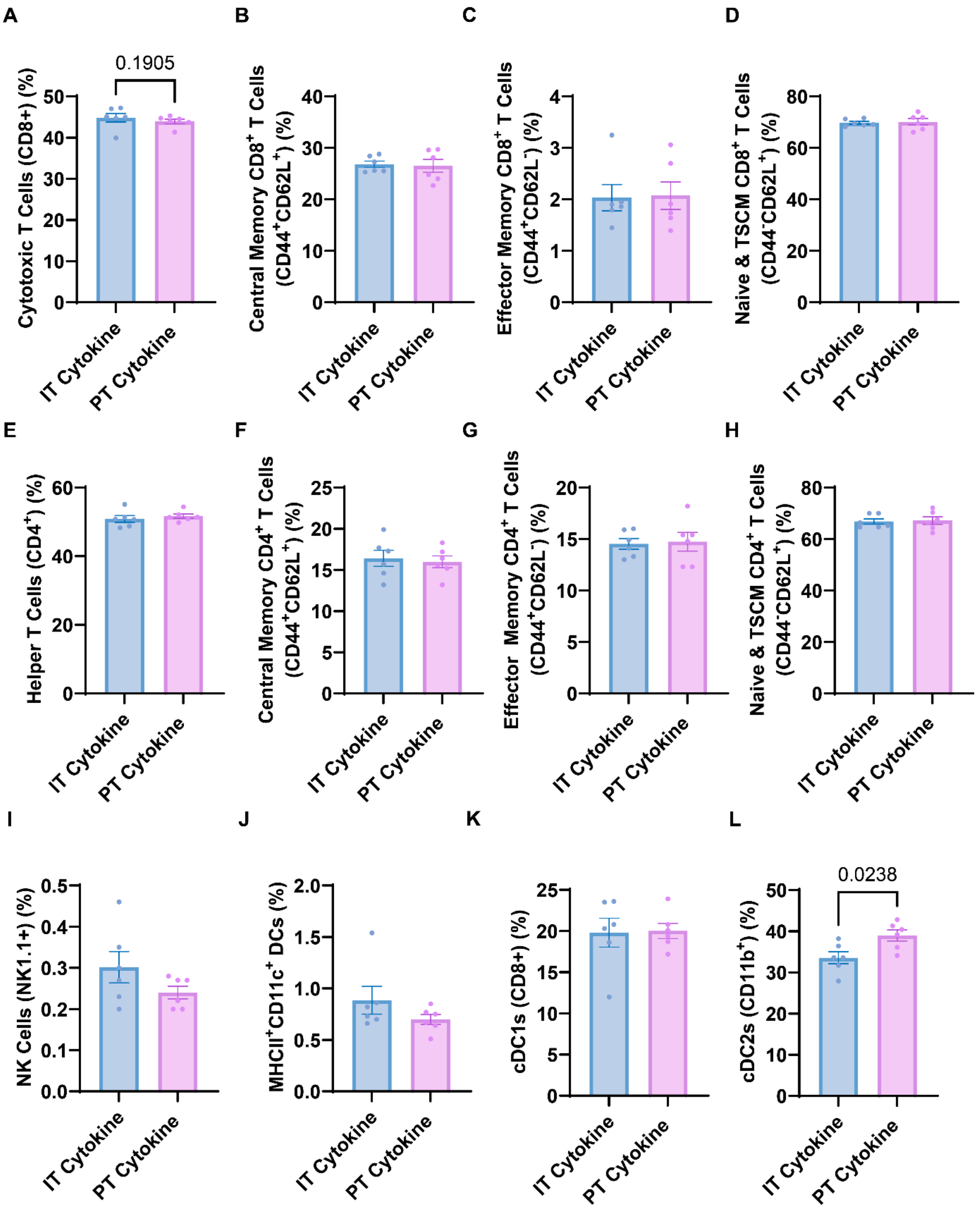


**Figure S5. Flow cytometry of lymph nodes on D7 post cytokine therapies. A.** Percent cytotoxic T cells (CD8$^+$) of CD3$^+$ T cells. **B.** Quantification of percentage of CD44$^+$CD62L$^+$ central memory T cells of CD8$^+$ T cells **C.** Quantification of CD44$^+$CD62L$^-$ effector memory T cells of CD8$^+$ T cells. **D.** Quantification of percentage of CD44$^-$CD62L$^+$ naïve and stem-like memory T cells of CD8$^+$ T cells. **E.** Percent cytotoxic T cells (CD4$^+$) of CD3$^+$ T cells. **F.** Quantification of percentage of CD44$^+$CD62L$^+$ central memory T cells of CD4$^+$ T cells **G.** Quantification of CD44$^+$CD62L$^-$ effector memory T cells of CD4$^+$ T cells. **H.**

Quantification of percentage of $CD44^{-}CD62L^{+}$ naïve and stem-like memory T cells of $CD4^{+}$ T cells. **I.** Quantification of percent of NK cells ($NK1.1^{+}$) cells of $CD45^{+}$ cells **J.** Percent of $MCHII^{+}$ $CD11c^{+}$ DCs of $CD45^{+}$ cells **K.** Percent of cDC1s ($CD8^{+}$) of $MCHII^{+}$ $CD11c^{+}$ DCs **L.** Percent of cDC2s ($CD11b^{+}$) of $MCHII^{+}$ $CD11c^{+}$ DCs. N=6 for all groups. Data reported as mean +/- SEM. Statistics determined by two-tailed Mann Whitney test in GraphPad prism.

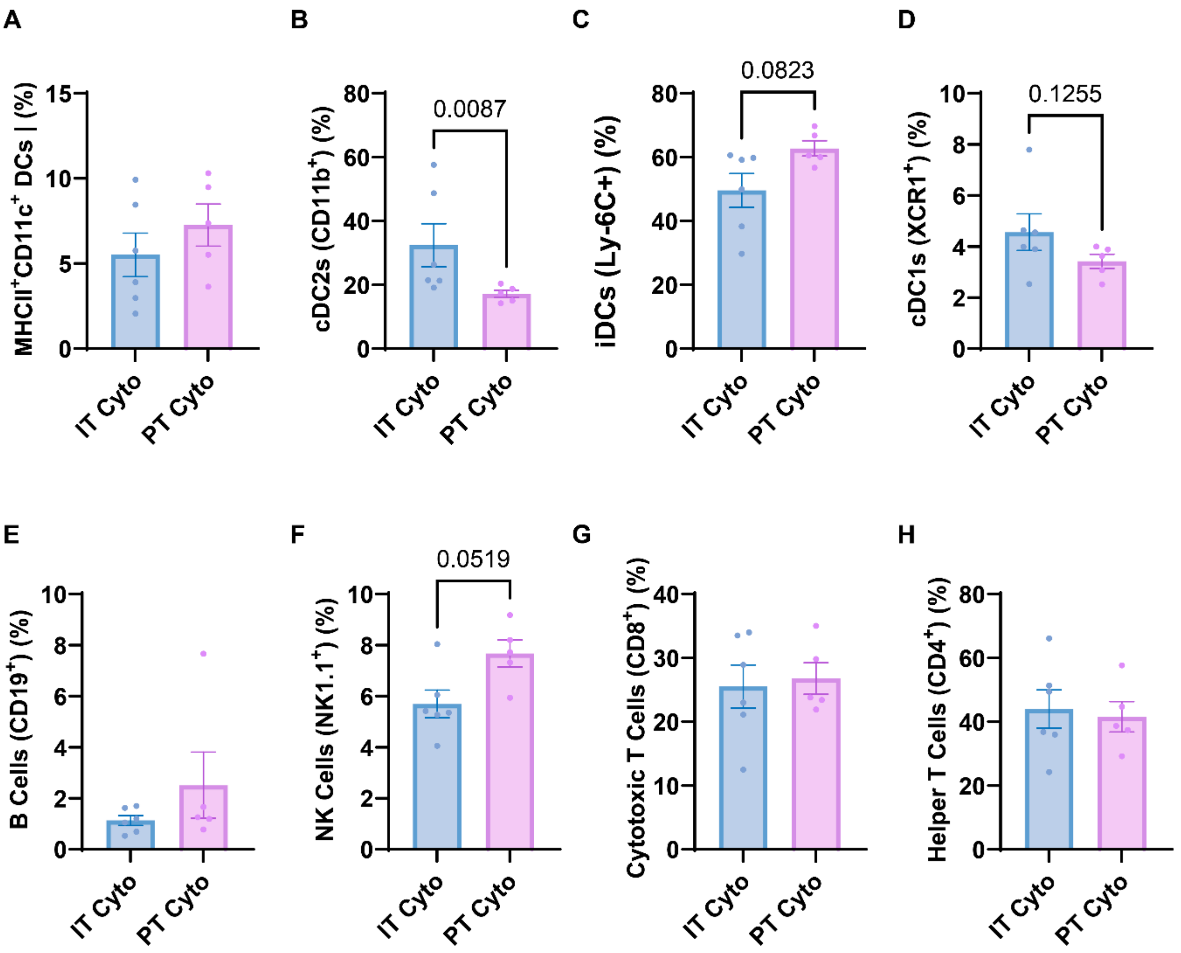


**Figure S6. Flow cytometry of tumors on day 3 post cytokine therapies. A.** Percent of MCHII$^+$ CD11c$^+$ DCs of CD45$^+$ cells **B.** Percent of cDC2s (CD11b$^+$) of MCHII$^+$ CD11c$^+$ DCs **C.** Percent of iDCs (Ly-6C$^+$) cells of MCHII$^+$ CD11c$^+$ DCs **D.** Percent cDC1s (XCR1$^+$) cells of MCHII$^+$ CD11c$^+$ DCs **E.** Percent of B cells (CD19$^+$) of CD45$^+$ cells **F.** Percent of NK cells (NK1.1$^+$) cells of CD45$^+$ cells **G.** Percent of CD8$^+$ T cells of CD3$^+$ T cells **H.** Percent of CD4$^+$ T cells of CD3$^+$ T cells. N=6 for IT cytokine, N=5 for PT cytokine. Data reported as mean +/- SEM. Statistics determined by two-tailed Mann Whitney test in GraphPad prism.

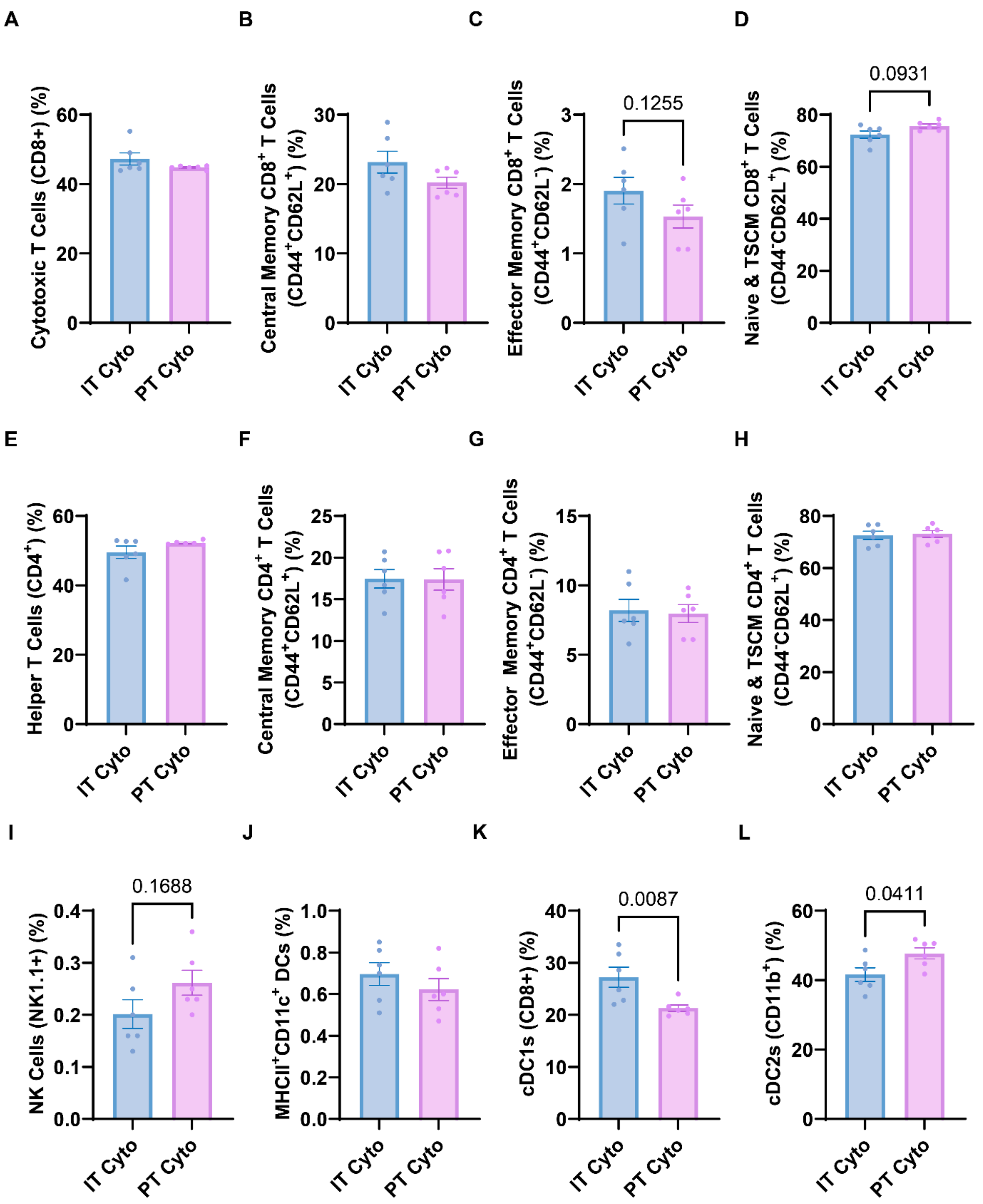


**Figure S7. Flow cytometry of lymph nodes on day 3 post cytokine therapies. A.** Percent cytotoxic T cells (CD8$^{+}$) of CD3$^{+}$ T cells. **B.** Quantification of percentage of CD44$^{+}$CD62L$^{+}$ central memory T cells of CD8$^{+}$ T cells **C.** Quantification of CD44$^{+}$CD62L$^{-}$ effector memory T cells of CD8$^{+}$ T cells. **D.** Quantification of percentage of CD44$^{-}$CD62L$^{+}$ naïve and stem-like memory T cells of CD8$^{+}$ T cells. **E.** Percent cytotoxic T cells (CD4$^{+}$) of CD3$^{+}$ T cells. **F.** Quantification of percentage of CD44$^{+}$CD62L$^{+}$ central memory T cells of CD4$^{+}$ T cells **G.** Quantification of CD44$^{+}$CD62L$^{-}$ effector memory T cells of CD4$^{+}$ T cells. **H.**

Quantification of percentage of $CD44^-CD62L^+$ naïve and stem-like memory T cells of $CD4^+$ T cells. **I.** Quantification of percent of NK cells ($NK1.1^+$) cells of $CD45^+$ cells **J.** Percent of $MCHII^+$ $CD11c^+$ DCs of $CD45^+$ cells **K.** Percent of cDC1s ($CD8^+$) of $MCHII^+$ $CD11c^+$ DCs **L.** Percent of cDC2s ($CD11b^+$) of $MCHII^+$ $CD11c^+$ DCs. N=6 for all groups. Data reported as mean +/- SEM. Statistics determined by two-tailed Mann Whitney test in GraphPad prism.

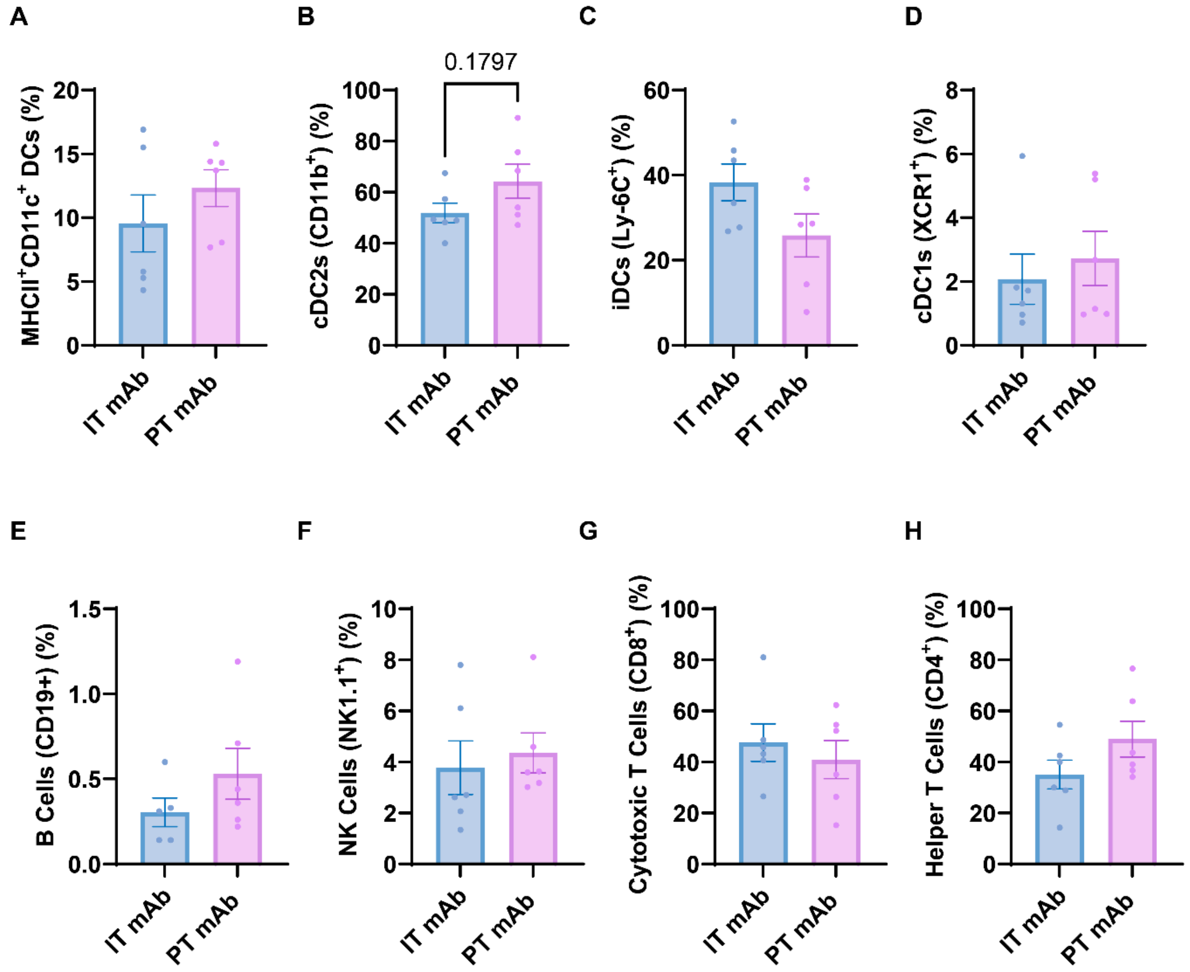


**Figure S8. Additional flow cytometry of tumors on day 7 post antibody therapies. A.** Percent of MCHII$^+$ CD11c$^+$ DCs of CD45$^+$ cells **B.** Percent of cDC2s (CD11b$^+$) of MCHII$^+$ CD11c$^+$ DCs **C.** Percent of iDCs (Ly-6C$^+$) cells of MCHII$^+$ CD11c$^+$ DCs **D.** Percent cDC1s (XCR1$^+$) cells of MCHII$^+$ CD11c$^+$ DCs **E.** Percent of B cells (CD19$^+$) of CD45$^+$ cells **F.** NK cells (NK1.1$^+$) cells of CD45$^+$ cells **G.** Percent of CD8$^+$ T cells of CD3$^+$ T cells **H.** Percent of CD4$^+$ T cells of CD3$^+$ T cells. N=6 for all groups. Data reported as mean +/- SEM. Statistics determined by two-tailed Mann Whitney test in GraphPad prism.

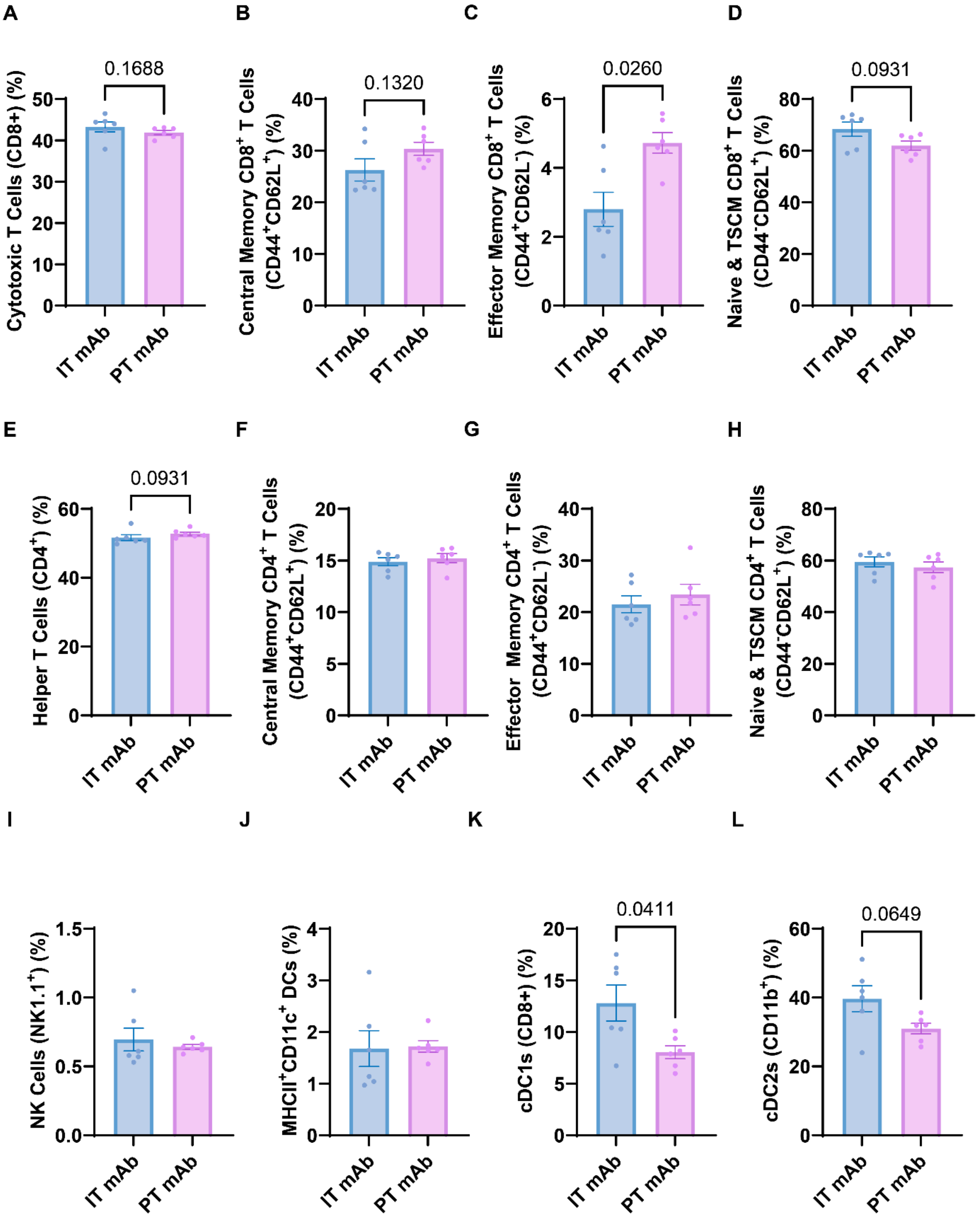


**Figure S9. Flow cytometry of lymph nodes day 7 post antibody therapies. A.** Percent cytotoxic T cells ($CD8^+$) of $CD3^+$ T cells. **B.** quantification of percentage of $CD44^+CD62L^+$ central memory T cells of $CD8^+$ T cells **C.** quantification of $CD44^+CD62L^-$ effector memory T cells of $CD8^+$ T cells. **D.** quantification of percentage of $CD44^-CD62L^+$ naïve and stem-like memory T cells of $CD8^+$ T cells. **E.** Percent cytotoxic T cells ($CD4^+$) of $CD3^+$ T cells. **F.** quantification of percentage of $CD44^+CD62L^+$ central memory T cells of

CD4$^+$ T cells **G.** quantification of CD44$^+$CD62L$^-$ effector memory T cells of CD4$^+$ T cells. **H.** quantification of percentage of CD44$^-$CD62L$^+$ naïve and stem-like memory T cells of CD4$^+$ T cells. **I.** quantification of percent of NK cells (NK1.1$^+$) cells of CD45$^+$ cells **J.** percent of MCHII$^+$ CD11c$^+$ DCs of CD45$^+$ cells **K.** percent of cDC1s (CD8$^+$) of MCHII$^+$ CD11c$^+$ DCs **L.** percent of cDC2s (CD11b$^+$) of MCHII$^+$ CD11c$^+$ DCs. N=6 for all groups. Data reported as mean +/- SEM. Statistics determined by two-tailed Mann Whitney test in GraphPad prism.

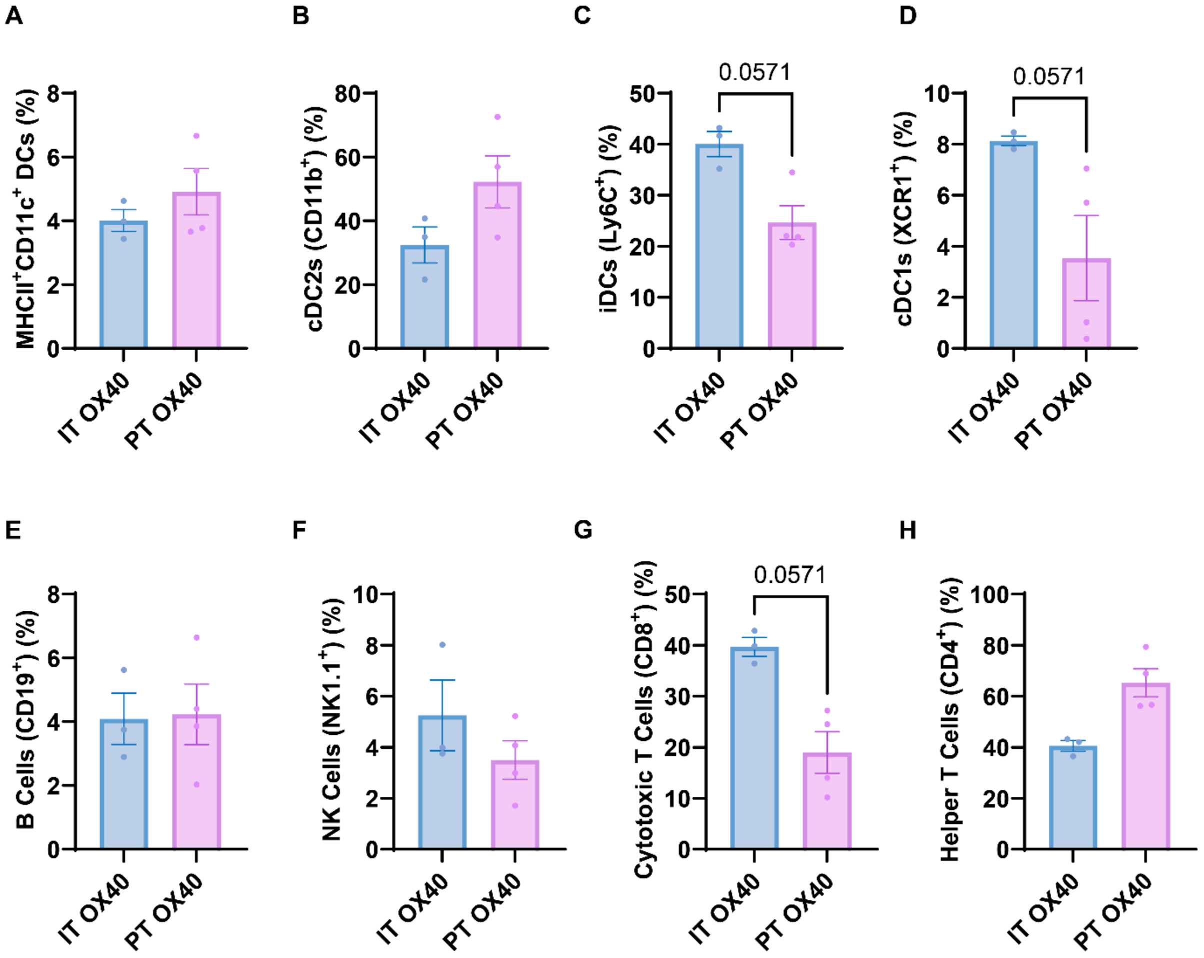


**Figure S10. Flow cytometry of tumors on day 14 post antibody therapies. A.** Percent of MCHII$^+$ CD11c$^+$ DCs of CD45$^+$ cells **B.** Percent of cDC2s (CD11b$^+$) of MCHII$^+$ CD11c$^+$ DCs **C.** Percent of iDCs (Ly-6C$^+$) cells of MCHII$^+$ CD11c$^+$ DCs **D.** Percent cDC1s (XCR1$^+$) cells of MCHII$^+$ CD11c$^+$ DCs **E.** Percent of B cells (CD19$^+$) of CD45$^+$ cells **F.** NK cells (NK1.1$^+$) cells of CD45$^+$ cells **G.** Percent of CD8$^+$ T cells of CD3$^+$ T cells **H.** Percent of CD4$^+$ T cells of CD3$^+$ T cells. N=3 for IT OX40a, N=4 for PT OX40a. Data reported as mean +/- SEM. Statistics determined by two-tailed Mann Whitney test in GraphPad prism.

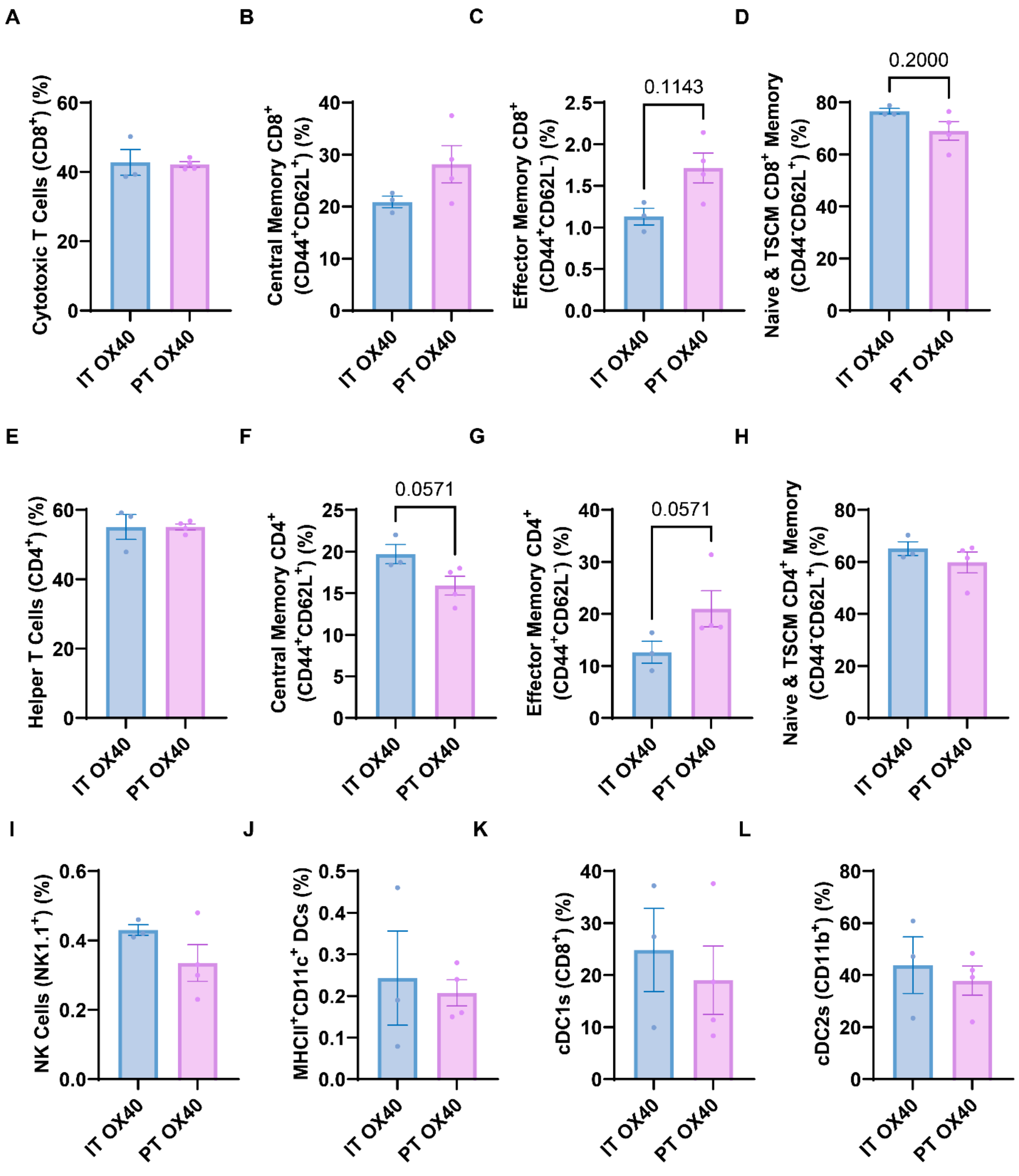


**Figure S11. Flow cytometry of lymph nodes on day 14 post antibody therapies. A.** Percent cytotoxic T cells ($CD8^+$) of $CD3^+$ T cells. **B.** Quantification of percentage of $CD44^+CD62L^+$ central memory T cells of $CD8^+$ T cells **C.** Quantification of $CD44^+CD62L^-$ effector memory T cells of $CD8^+$ T cells. **D.** Quantification of percentage of $CD44^-CD62L^+$ naïve and stem-like memory T cells of $CD8^+$ T cells. **E.** Percent cytotoxic T cells ($CD4^+$) of $CD3^+$ T cells. **F.** Quantification of percentage of $CD44^+CD62L^+$ central memory T cells of $CD4^+$ T cells **G.** Quantification of $CD44^+CD62L^-$ effector memory T cells of $CD4^+$ T cells. **H.** Quantification of percentage of $CD44^-CD62L^+$ naïve and stem-like memory T cells of $CD4^+$ T cells. **I.** Quantification of percent of NK cells ($NK1.1^+$) cells of $CD45^+$ cells **J.** Percent of $MCHII^+$ $CD11c^+$ DCs of

CD45$^+$ cells **K.** Percent of cDC1s (CD8$^+$) of MCHII$^+$ CD11c$^+$ DCs **L.** Percent of cDC2s (CD11b$^+$) of MCHII$^+$ CD11c$^+$ DCs. N=3 for IT OX40a, N=4 for PT OX40a. Data reported as mean +/- SEM. Statistics determined by two-tailed Mann Whitney test in GraphPad prism.

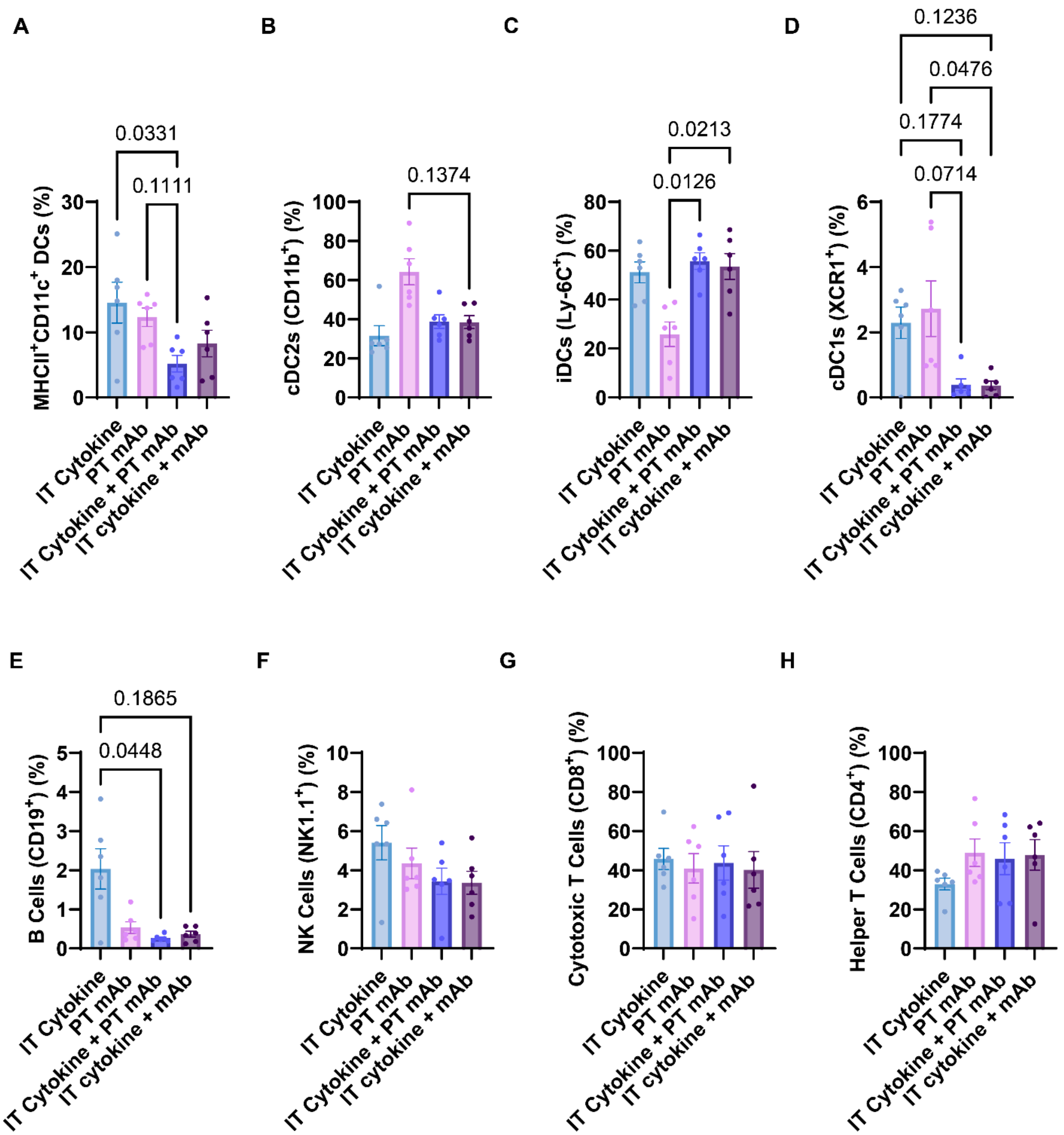


**Figure S12. Flow cytometry of tumors on day 7 post combination therapies. A.** Percent of MCHII$^+$ CD11c$^+$ DCs of CD45$^+$ cells **B.** Percent of cDC2s (CD11b$^+$) of MCHII$^+$ CD11c$^+$ DCs **C.** Percent of iDCs (Ly-6C$^+$) cells of MCHII$^+$ CD11c$^+$ DCs **D.** Percent cDC1s (XCR1$^+$) cells of MCHII$^+$ CD11c$^+$ DCs **E.** Percent of B cells (CD19$^+$) of CD45$^+$ cells **F.** NK cells (NK1.1$^+$) cells of CD45$^+$ cells **G.** Percent of CD8$^+$ T cells of CD3$^+$ T cells **H.** Percent of CD4$^+$ T cells of CD3$^+$ T cells. N=6 for all groups. Data reported as mean +/- SEM. Statistics determined by Kruskal-Wallis test with Dunn's multiple comparisons test in GraphPad prism.
